# Transcriptomic profiling of *Debaryomyces hansenii* reveals detoxification and stress responses to benzo(a)pyrene exposure

**DOI:** 10.1101/2025.06.17.660189

**Authors:** Francisco Padilla-Garfias, Augusto César Poot-Hernández, Minerva Araiza-Villanueva, Martha Calahorra, Norma Silvia Sánchez, Antonio Peña

## Abstract

The environmental accumulation of polycyclic aromatic hydrocarbons (PAHs), such as benzo(a)pyrene (BaP), poses significant threats to ecosystems and public health due to their persistent nature, mutagenic potential, and well-documented carcinogenicity. In this study, we investigated the ability of the extremophilic yeast *Debaryomyces hansenii* to activate specialized detoxification mechanisms for BaP degradation, even under nutrient-deprived conditions. When exposed to 100 ppm BaP, *D. hansenii* eliminated over 70% of the contaminant within three days while maintaining normal growth dynamics. RNA-Seq analysis revealed widespread transcriptional remodeling, with 1179 genes upregulated and 1031 downregulated under BaP-only conditions, and 1067 upregulated and 977 downregulated genes during co-metabolic exposure (2% glucose + 100 ppm BaP), from a total of 6506 annotated genes. Gene Ontology (GO) and KEGG enrichment analyses highlighted the activation of xenobiotic degradation pathways, notably involving cytochrome P450 monooxygenases (CYPs), epoxide hydrolases (EHs), and glutathione S-transferases (GSTs), alongside an enhanced antioxidant response and finely tuned glutathione homeostasis. This work provides the first comprehensive transcriptomic profile of BaP detoxification in *D. hansenii*, revealing an intricate and highly adaptive stress response. Collectively, these findings position *D. hansenii* as a promising eukaryotic platform for bioremediation in saline and contaminated environments, especially where conventional microbial candidates fall short due to environmental extremes or nutrient scarcity.

**Importance:** Polycyclic aromatic hydrocarbons (PAHs), such as benzo(a)pyrene (BaP), are long-lasting environmental pollutants with serious health and ecological implications. Although microbial degradation offers a promising strategy for remediation, most efforts have focused on bacterial and filamentous fungal systems, leaving other microbial groups comparatively unexplored. In contrast, extremotolerant yeasts remain largely overlooked despite their inherent resilience. Here, we investigated the marine yeast *Debaryomyces hansenii* and discovered that it not only tolerates BaP under nutrient-limited conditions but also actively degrades it. This response relies on a combination of detoxifying enzymes and antioxidant defenses, reflecting a well-orchestrated metabolic adaptation to chemical stress. Our findings highlight the untapped and promising potential of *D. hansenii* as a robust eukaryotic chassis for bioremediation, particularly in environments where conventional microbes may fail to survive.

## 1. Introduction

Benzo(a)pyrene (BaP) is a five-ring polycyclic aromatic hydrocarbon (PAH) commonly produced during the incomplete combustion of organic matter and found in coal processing residues, oil sludge, asphalt, and tobacco smoke (1). Its remarkable chemical stability, conferred by its aromatic ring structure, underlies its environmental persistence and widespread distribution in soil and airborne particles (2–4). Human exposure is common, primarily through inhalation or dietary intake. Once absorbed, BaP is rapidly metabolized by cytochrome P450 (CYP) enzymes into reactive and often carcinogenic intermediates (1, 5, 6). These metabolites have been linked to a range of harmful effects, including epigenetic disruption, neurotoxicity, teratogenicity, and impaired reproductive function (6, 7).

One of the most promising strategies for mitigating BaP contamination is mycoremediation, that is a fungal-based approach that leverages the metabolic capacity of fungi to degrade persistent organic pollutants (8, 9). Fungal species capable of metabolizing PAHs are mainly found within the Ascomycota and Basidiomycota phyla and include both filamentous fungi and yeasts (9–12). However, most studies to date have focused on filamentous forms, using biochemical assays to evaluate degradation kinetics, leaving the potential of yeasts comparatively underexplored (13, 14).

Fungal degradation of PAHs involves a sequence of extracellular and intracellular reactions generally grouped into three metabolic phases. In Ascomycota, phase I the oxidative transformation of PAH into hydroxylated or epoxide intermediates, primarily mediated by cytochrome P450 monooxygenases (CYPs) and epoxide hydrolases (EHs). Phase II involves the conjugation of these reactive metabolites with sulfate, glucose, or reduced glutathione (GSH), catalyzed by transferases, such as glutathione S-transferases (GSTs). Finally, phase III facilitates the compartmentalization or excretion of the resulting water-soluble conjugates (10, 11, 15–17). These enzymatic processes are governed by a multigene regulatory system (collectively termed the xenome) that orchestrates fungal xenobiotic metabolism (10, 11, 18).

Co-metabolism, wherein a readily metabolizable carbon source is supplied alongside a recalcitrant compound, can markedly enhance PAH degradation by providing not only additional energy but also key cofactors for detoxification enzyme activity. While BaP degradation is the primary focus of this study, we also examined how glucose supplementation might influence this process. Previous studies have shown that glucose availability can stimulate the activity of key detoxification enzymes, including CYPs, EHs, and GSTs, thereby supporting both growth and detoxification (19, 20). However, how such modulation plays out in extremophilic yeasts like *D. hansenii* remains largely uncharacterized.

Among non-conventional yeasts, *Debaryomyces hansenii* has emerged as a particularly versatile and stress-tolerant species, capable of thriving under high salinity, oxidative stress, low temperatures, and even in the presence of heavy metals and hydrophobic hydrocarbons such as BaP (21–23). Its broad environmental adaptability, combined with its oleaginous metabolism and metabolic plasticity, makes it a compelling candidate for bioremediation applications, especially in extreme or contaminated ecosystems (24–27).

While several yeasts, such as *Candida*, *Cryptococcus*, *Rhodotorula*, *Pichia*, and *Debaryomyces* spp., are known to degrade high molecular weight PAHs (10), only *Rhodotorula mucilaginosa* has been analyzed at the transcriptomic level using next-generation sequencing (NGS) (28). These studies revealed the transcriptional activation of detoxification, redox balance, and DNA repair pathways, emphasizing the central role of these mechanisms in enabling yeast survival under xenobiotic stress(10, 28, 29).

However, to date, no transcriptomic studies have explored how extremophilic yeasts like *D. hansenii* respond to BaP or similar PAHs. Most available transcriptomic data instead focus on halotolerance and metal resistance (27, 30). Interestingly, the presence of a functional CYP gene (*DhDIT2*) capable of BaP oxidation adds further weight to its potential as a biotechnological chassis for use in marine or saline environments (23, 25). In this study, we present the first genome-wide transcriptomic analysis of *D. hansenii* exposed to BaP, under both nutrient-limited and co-metabolic conditions. By integrating RNA-Seq with physiological assays, we aim to elucidate the molecular principles of BaP detoxification and showcase the promise of *D. hansenii* as an extremotolerant platform for sustainable mycoremediation.

## 2. Materials and methods

### 2.1 Strains, media, and culture conditions

*Debaryomyces hansenii* Y7426 strain (United States Department of Agriculture, Peoria, Illinois, USA) was maintained on solid YNBG medium (0.67% yeast nitrogen base with amino acids, ammonium sulphate, supplemented with 2% glucose, and 2% agar). Cultures were refreshed monthly and routinely grown overnight in 250 mL of YNBG within 500 mL Erlenmeyer flasks at 28 °C with orbital shaking at 250 revolutions per minute (rpm) for 24 hours.

A 10,000 ppm (10 mg/mL) stock solution of BaP (Sigma-Aldrich B1760) was prepared by dissolving 10 mg of BaP in 1.0 mL of analytical-grade acetone. The stock solution was added to culture medium to reach a final concentration of 100 ppm (100 µg/mL or ∼396 µM). The final acetone concentration was kept below 1% (v/v), a condition previously shown not to affect the growth or metabolic activity of *D. hansenii* (23).

### 2.2 Growth and survival assays

All growth and survival assays were performed under aseptic conditions using three independent biological replicates, each measured in technical duplicate, unless otherwise indicated. To assess BaP tolerance, *D. hansenii* cells were pre-cultured for 24 hours in YNBG, washed twice with sterile water, and resuspended to an initial optical density at 600 nm (OD_600_) of 1.0. Serial 10-fold dilutions were prepared in sterile water in 96-well plates, and 5 µL of each dilution were spotted onto the surface of YNB, YNBG, YNB + 100 ppm BaP, or YNBG + 100 ppm BaP plates, using an aluminum multi-pin replicator. Plates were incubated at 28 °C and monitored daily for six days.

For growth curves analysis, 24-hour pre-culture was adjusted to an OD_600_ of 0.03, using a Beckman DU 650 spectrophotometer and inoculated into the respective media. Cultures were incubated at 28 °C, and optical density was recorded hourly for six days using a Bioscreen C automated plate reader (with an initial OD_600_ baseline of approximately 0.2 in this instrument).

Dry weight determinations were performed over 6 days using the method described by Peña *et al*. (2023) (31). Pre-cultures grown in YNBG were used to inoculate 50 mL of fresh YNB, YNBG, YNB + 100 ppm BaP, or YNBG + 100 ppm BaP, adjusting to OD_600_ = 0.1. At 24 hours intervals, 1.0 mL aliquots were collected, transferred to aluminum plates, and dried at 95 °C until a constant weight.

### 2.3 BaP degradation assay

BaP degradation was quantified by spectrofluorometry and validated by gas chromatography-mass spectrometry (GC-MS), following the method described by Padilla-Garfias *et al*. (2022) (23). All assays were conducted in triplicate using three independent biological replicates. Cultures of *D. hansenii* grown for 24 hours in YNBG were used to inoculate 250 mL of YNB or YNBG supplemented with 100 ppm BaP, adjusted to an initial OD_600_ = 0.1. Samples (3.0 mL) were collected at 0, 1, 2, 3, and 6 days and stored at −20 °C for BaP extraction.

Extraction was performed with chloroform, followed by evaporation and resuspension in acetone. Fluorescence was measured using an AMINCO SLM spectrofluorometer (excitation at 356 nm, emission at 405 nm), and concentrations were calculated from a standard curve. To distinguish true biodegradation from passive adsorption or abiotic loss, two negative controls were included: a cell-free medium control and a heat-inactivated culture control (95 °C, 15 minutes).

GC-MS analysis was conducted using a Hewlett-Packard 5890-II system with a JEOL SX-102 A spectrometer and an HP-5MS column (30 m × 0.25 mm, 0.25 μm). Helium served as the carrier gas; chromatographic separation used a temperature gradient from 50 °C to 300 °C. Detection parameters followed standard ionization protocols.

### 2.4 Glucose consumption

Glucose consumption was measured using the Glucose Oxidase Activity Assay Kit MAK501 (Sigma-Aldrich, USA), with slight modifications to the manufacturer’s protocol. *D. hansenii* was cultured in 50 mL of YNBG or YNBG supplemented with 100 ppm BaP in 150 mL Erlenmeyer flasks at 28 °C with shaking at 250 rpm for 6 days. Each condition was evaluated using three independent biological replicates, with technical duplicates per sample.

Aliquots (1.0 mL) were collected at 0, 1, 2, 3, and 6 days, centrifuged (3100 rpm, 1625 × g), and the supernatants were stored at −20 °C. Before analysis, samples were thawed and diluted 1:10 in deionized water. A 50 µL aliquot was mixed with 100 µL of Glucose Oxidase Reagent and 50 µL of the colorimetric substrate, incubated at room temperature for 30 minutes, and absorbance was measured at 570 nm.

Glucose concentration was determined using a standard curve (0–10 mM), and consumption was calculated as the difference between the initial value (day 0) and subsequent measurements.

### 2.5 RNA Extraction

*D. hansenii* cells were initially grown in 250 mL of YNBG medium at 28 °C with shaking at 250 rpm for 24 hours. Following this pre-culture, two separate RNA-Seq experiments were performed using three independent biological replicates per condition. In the first, cells were transferred to 50 mL flasks containing either YNBG or YNB supplemented with 100 ppm BaP and incubated under the same conditions for 3 days. In the second experiment, cells were cultured in either YNBG or YNBG + 100 ppm BaP.

For RNA extraction, the same protocol was applied to all samples across the study. A total of 15 mL of culture was collected on day 0 (post pre-culture) and day 3 (in all conditions). Cells were pelleted by centrifugation and resuspended in 1.0 mL of AE buffer (50 mM sodium acetate, 10 mM EDTA, pH 5.2). Total RNA was extracted using a modification of the method described by Schmitt *et al*. (1990) (32). RNA quality and integrity were assessed by 1% denaturing agarose gel electrophoresis, confirming the presence of intact 28S and 18S rRNA bands.

### 2.6 RNA-Seq: sample preparation, library construction, data analysis, gene enrichment

Total RNA was purified from samples collected and extracted as described in Section 2.5, using the same standardized protocol across all conditions. RNA was precipitated with 7.5 M LiCl following the method of Walker *et al*. (2013) (33). RNA quality and integrity were verified through 1% denaturing agarose gel electrophoresis, spectrophotometry (NanoDrop, Thermo Scientific), and Bioanalyzer 2100 (Agilent Technologies). Only samples with OD260/280 and OD260/230 ratios above 2.0, and RIN values ≥4.0, were included for sequencing. Three independent biological replicates per condition were analyzed.

RNA samples were lyophilized and sent to Novogene Co. (Sacramento, CA, USA) for library preparation and sequencing. Poly(A)-enriched mRNA was isolated using magnetic beads with oligo(dT), fragmented, and reverse-transcribed using random hexamer primers. Library construction involved end repair, adapter ligation, size selection (∼150 bp), PCR amplification, and quality control via Qubit and RT-PCR. Paired-end sequencing (≥20 million read pairs per sample) was performed on the Illumina HiSeq 6000 platform. Bioinformatic processing was conducted using the snakePipes (v2.5.0) mRNA-seq pipeline (34). In more detail, raw reads and mapping were quality-checked with FastQC and MultiQC (35), adapters were trimmed with Cutadapt (36), and sequences were aligned to the *D. hansenii* CBS767 genome (NCBI GCF_000006445.2) using RNA STAR (37). Read counts per gene were obtained with FeatureCounts (38) and differential expression was analyzed using DESeq2 (39).

Functional enrichment of differentially expressed genes was performed using FungiFun (40), based on Gene Ontology (GO) terms (41, 42) and Kyoto Encyclopedia of Genes and Genomes (KEGG) annotations (43–45) Enrichment was considered significant at an adjusted p-value < 0.05. KEGG pathway visualization was conducted via the KEGG REST-api (43–45).

### 2.7 RNA-seq verification: cDNA synthesis and RT-qPCR to analyze gene expression

To validate the RNA-seq results and perform gene expression experiments, total RNA was extracted as described in Section 2.5, treated with DNase I (RQ1 RNase-Free DNase kit, Promega) to eliminate genomic DNA contamination, and reverse transcribed into cDNA using the ImProm-II™ Reverse Transcription System (Promega). Gene expression validation was subsequently performed by RT-qPCR using three independent biological replicates and two technical replicates per sample.

RT-qPCR was conducted using specific primers (see Supplementary Table 1) targeting the 10 most up- and downregulated open reading frames (ORFs), as well as genes associated with detoxification pathways. Sequences were retrieved from NCBI, and primers were designed using Primer-BLAST, then evaluated for dimer formation and specificity with DINAMelt (http://www.unafold.org/hybrid2.php) (46). Primers were synthesized at the Molecular Biology Unit of the Institute of Cellular Physiology, UNAM.

Reactions were run on a Rotor-Gene Q thermal cycler (Qiagen) using SYBR Green chemistry (qPCR SyberMaster highROX, Jena Bioscience). Relative gene expression was quantified using the ΔΔCt method (47), with YNBG-grown cells as the calibrator. Expression values were normalized to *DhACT1* (ORF DEHA2D05412g), following Sánchez *et al*. (2020) (48).

### 2.8 Statistical analysis

Data are presented as mean ± standard deviation from three independent biological replicates, each measured in triplicate. Statistical significance was evaluated by two-way ANOVA in Prism 9 (GraphPad Software Inc., San Diego, CA, USA), with a 95% confidence interval. Differences were considered statistically significant at *p* < 0.05, and significance levels are indicated in the corresponding figure legends, for Figure 1, areas under the curve (AUC) were calculated to perform the aforementioned analyses, for more details see Supplementary Figure 1.

**Figure 1.**
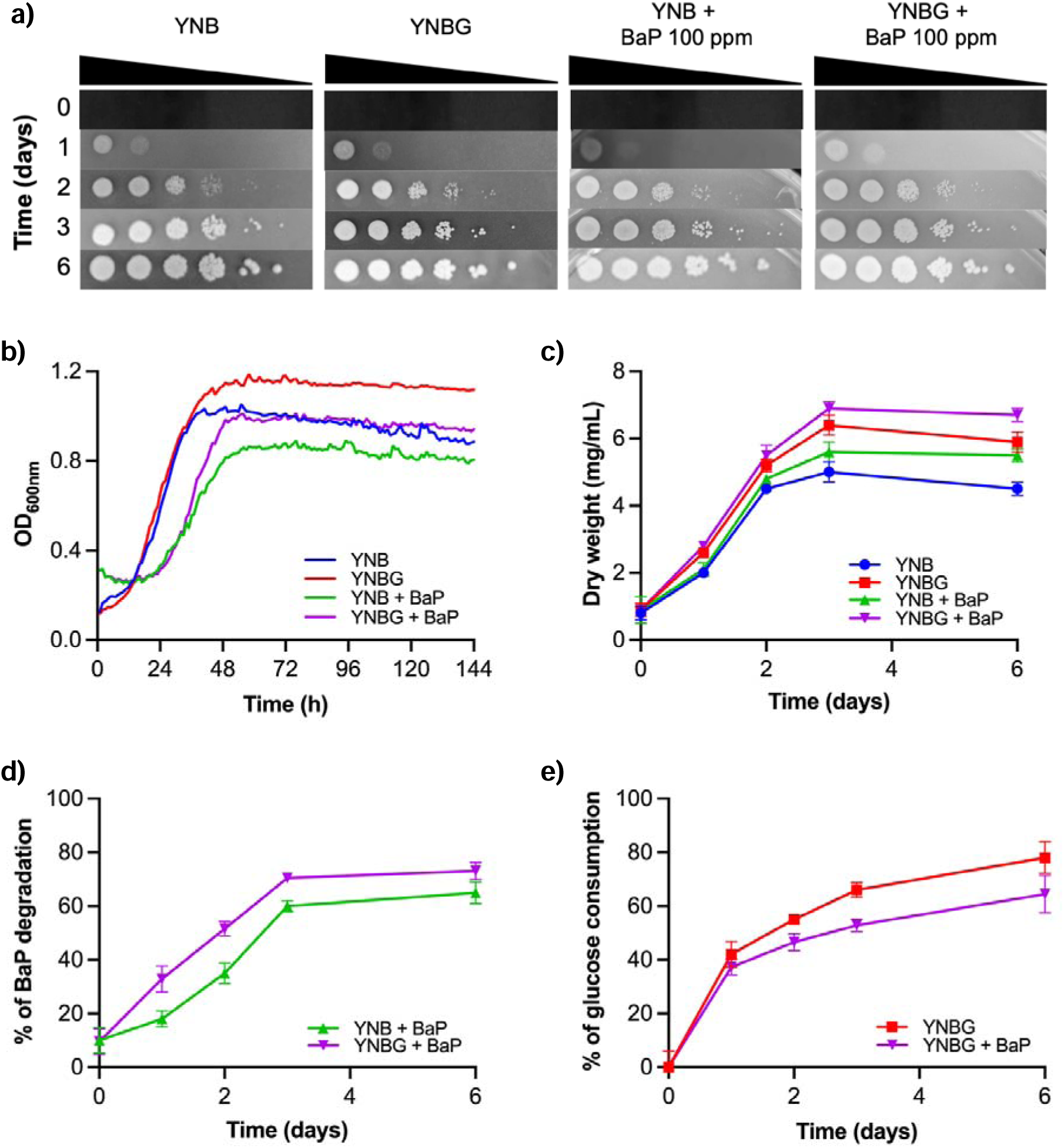
Physiological response of *D. hansenii* to BaP exposure. a) Growth on solid YNB, YNBG, YNB + 100 ppm BaP, and YNBG + 100 ppm BaP media assessed by serial dilution over 6 days. b) Growth curves in liquid cultures under the same conditions. c) Biomass quantification via dry weight measurements over time. d) BaP degradation kinetics evaluated by spectrofluorometry. e) Glucose consumption dynamics in the presence or absence of BaP. Statistical comparisons are provided in Supplementary Figure 1.

### 2.9 Data availability

The raw and processed RNA-Seq data generated in this study are available from the NCBI Gene Expression Omnibus database under acession No. GSE299919.

## 3. Results

Based on previous findings demonstrating *D. hansenii*’s ability to tolerate up to 500 ppm BaP and identifying *DhDIT2* as a BaP-oxidizing CYP gene (23), we performed a comprehensive set of physiological assays to assess yeast growth and metabolism under both nutrient-limited and co-metabolic conditions. These assays included growth evaluations on solid and liquid media, dry biomass measurements, BaP degradation kinetics, and glucose consumption analysis (Figure 1). Remarkably, spot assays confirmed that *D. hansenii* tolerates 100 ppm BaP as a sole carbon source, exhibiting growth patterns that closely resembled those observed under both control (YNBG) and minimal (YNB) conditions (Figure 1a).

Growth curve analysis (Figure 1b) revealed that *D. hansenii* consistently reached the stationary phase within 48 hours (2 days) across all experimental conditions, although a noticeable increase in doubling time revealed a clear metabolic burden imposed by BaP. In minimal medium, the generation time was notably higher (YNB: 10.0 ± 0.04 h) compared to glucose-supplemented culture (YNBG: 7.5 ± 0.02 h), and exposure to BaP further increased doubling times (YNB + BaP: 18.3 ± 2.7 h; YNBG + BaP: 11.7 ± 1.5 h).

It is important to mention and consider that the growth curves (Figure 1b) were performed in an automated plate reader where agitation and oxygen availability is not the same as in the experimental conditions in which dry weight (Figure 1c), BaP degradation (Figure 1d) and glucose consumption (Figure 1e) were determined. It is also important to note that the experimental strategy of Figures 1c–e was used for RNA extraction.

Dry weight measurements (Figure 1c) corroborated these observations. As expected, cultures exposed to BaP as the sole carbon source exhibited reduced biomass accumulation compared to control conditions. However, co-metabolic cultures (YNBG + BaP) produced the highest biomass levels, suggesting that *D. hansenii* can partially metabolize BaP to sustain growth, even under nutrient-limited (oligotrophic) conditions.

BaP degradation efficiency, assessed by spectrofluorometry and confirmed by GC-MS on day 6 of the degradation curve (Supplementary Figure 2), was notably higher under co-metabolic conditions (73.1%) compared to BaP-only cultures (65.0%). In both cases, degradation rates reached by day 3, indicating early stabilization of the detoxification process (Figure 1d). Importantly, the BaP recovery rate was 90% in a cell-free culture medium containing 100 ppm BaP, while the recovery rate was 87% in a medium inoculated with dead yeast cells (dead control), which was quantified as a control over the 6 days (data not shown).

Finally, glucose consumption profiles (Figure 1e) indicated that BaP exposure attenuated glucose uptake, with BaP-treated cultures consuming only 64.5% of available glucose versus 78% in controls by day 6. This reduction suggests a metabolic shift, potentially reflecting partial reliance on BaP as an alternative carbon or energy source.

Since BaP degradation stabilized and cultures entered the stationary phase by 72 hours (Figure 1), we selected this time point for RNA extraction and transcriptomic profiling. Quality control of cDNA libraries confirmed the expected fragment sizes (∼150 bp), and sequencing generated an average of 47.2 million paired-end reads per sample, with alignment rates consistently exceeding 95%. Approximately 45% of total reads mapped to annotated genes, a proportion comparable to previous reports in yeast transcriptomic studies (27).

Principal component analysis (PCA) of RNA-Seq data (Figure 2a) revealed a clear separation between BaP-exposed samples and controls, with BaP-treated replicates forming a tight cluster, indicative of a robust and consistent transcriptional response. In contrast, control replicates exhibited greater dispersion, suggesting a broader transcriptional variability under nutrient-supplemented conditions.

**Figure 2.**
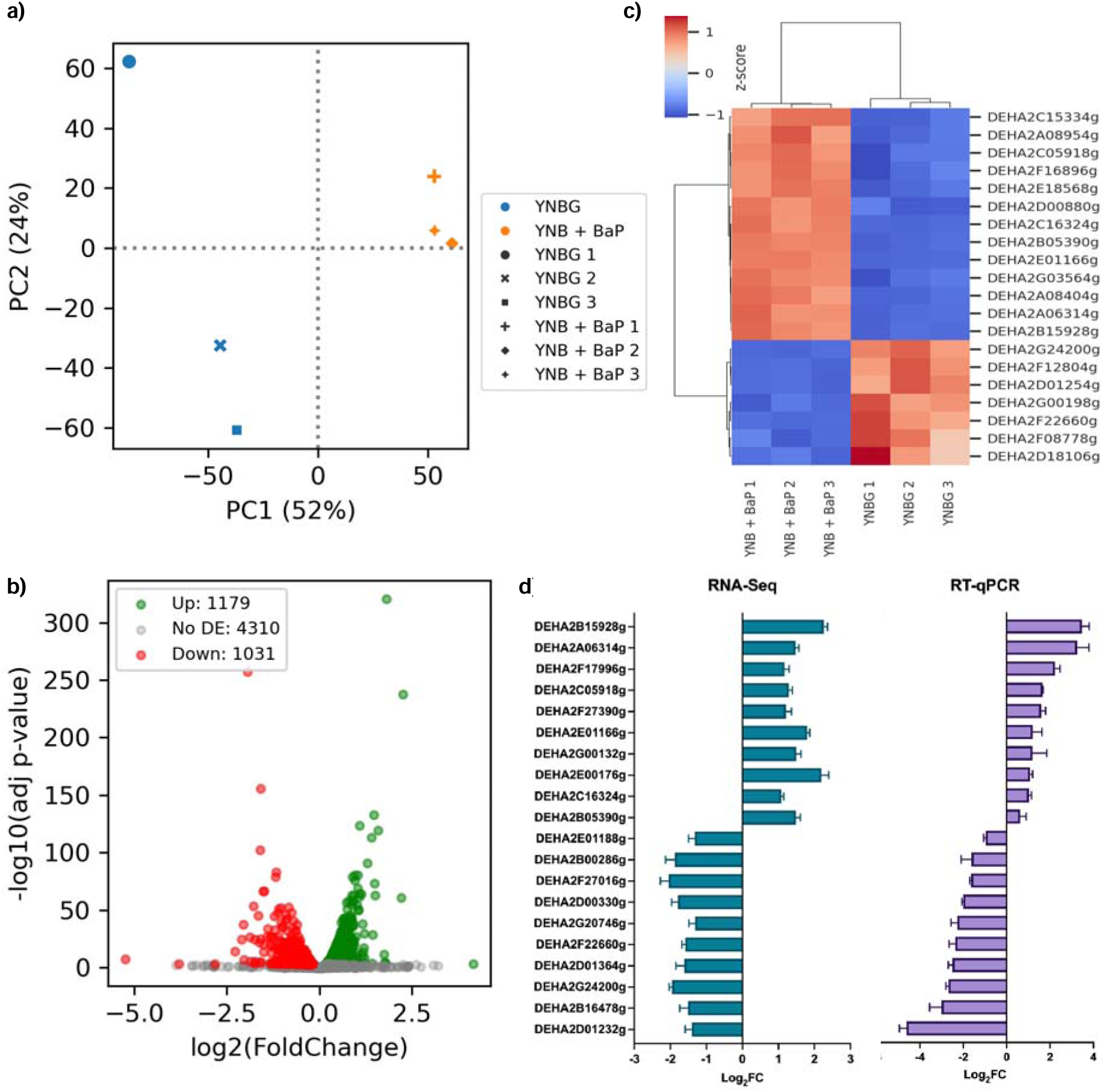
Transcriptomic analysis of *D. hansenii* exposed to 100 ppm of BaP. a) Principal component analysis. YNBG control (YNBG 1, YNBG 2, and YNBG 3) and YNB + BaP treatment (YNB + BaP 1, YNB + BaP 2, and YNB + BaP 3). b) Volcano plot obtained from the differential expression analysis (adjusted *p*-value < 0.05). c) Top 20 differentially expressed ORFs, upregulated (red) and downregulated (blue) genes based on RNA-Seq differential expression analysis (control condition: YNBG; experimental condition: YNB + BaP). d) RT-qPCR validation of the 10 most overexpressed and the 10 most underexpressed genes identified through RNA-Seq differential expression analysis.

Differential expression analysis identified 1179 upregulated and 1031 downregulated genes out of the 6506 annotated ORFs in BaP-treated cultures compared to control condition (Figure 2b). These transcriptional changes reflect a widespread and coordinated reprogramming of cellular processes in response to BaP.

The 20 most significant differentially expressed ORFs, both up- and downregulated, under BaP exposure are summarized in Figure 2c. Upregulated genes were associated with diverse biological processes, including glyoxalase detoxification (DEHA2D00880g), acetate transport (DEHA2G03564g), epoxide hydration (DEHA2A08404g), membrane structure (DEHA2B15928g), amino acid metabolism (DEHA2A06314g), GABA transamination (DEHA2C16324g), and various membrane transporters (DEHA2E01166g, DEHA2A08954g, DEHA2C05918g). Additional upregulated ORFs encoded methyltransferases (DEHA2B05390g), enzymes for carbohydrate storage (DEHA2E18568g), and several proteins of unknown function (e.g., DEHA2C15334g, DEHA2F16896g). Conversely, downregulated genes included those involved in calcium transport (DEHA2F12804g), protein folding (DEHA2F22660g), antibiotic response (DEHA2G00198g), and uncharacterized proteins (e.g., DEHA2G24200g, DEHA2D01254g, DEHA2F08778g, DEHA2D18106g). Functional annotations were retrieved from InterPro (49).

To validate the RNA-Seq findings, we performed RT-qPCR on the 10 most upregulated and 10 most downregulated ORFs (Figure 2d). The RT-qPCR results closely mirrored the RNA-Seq data in both the direction and magnitude of gene expression changes, confirming the reliability and reproducibility of the transcriptomic dataset (designed oligonucleotides Supplementary Table1).

Following differential gene expression analysis, we performed functional enrichment using Gene Ontology (GO) and Kyoto Encyclopedia of Genes and Genomes (KEGG) annotations to classify the upregulated genes (Figures 3a and 3b). Among the 1179 upregulated ORFs, 587 genes exhibited significant enrichment in GO terms related to the citric acid cycle, carbohydrate metabolism, redox regulation, membrane transport, and DNA repair. Several additional genes encoded proteins implicated in preserving membrane integrity, collectively suggesting a coordinated and multifaceted cellular adaptation to BaP-induced stress.

**Figure 3.**
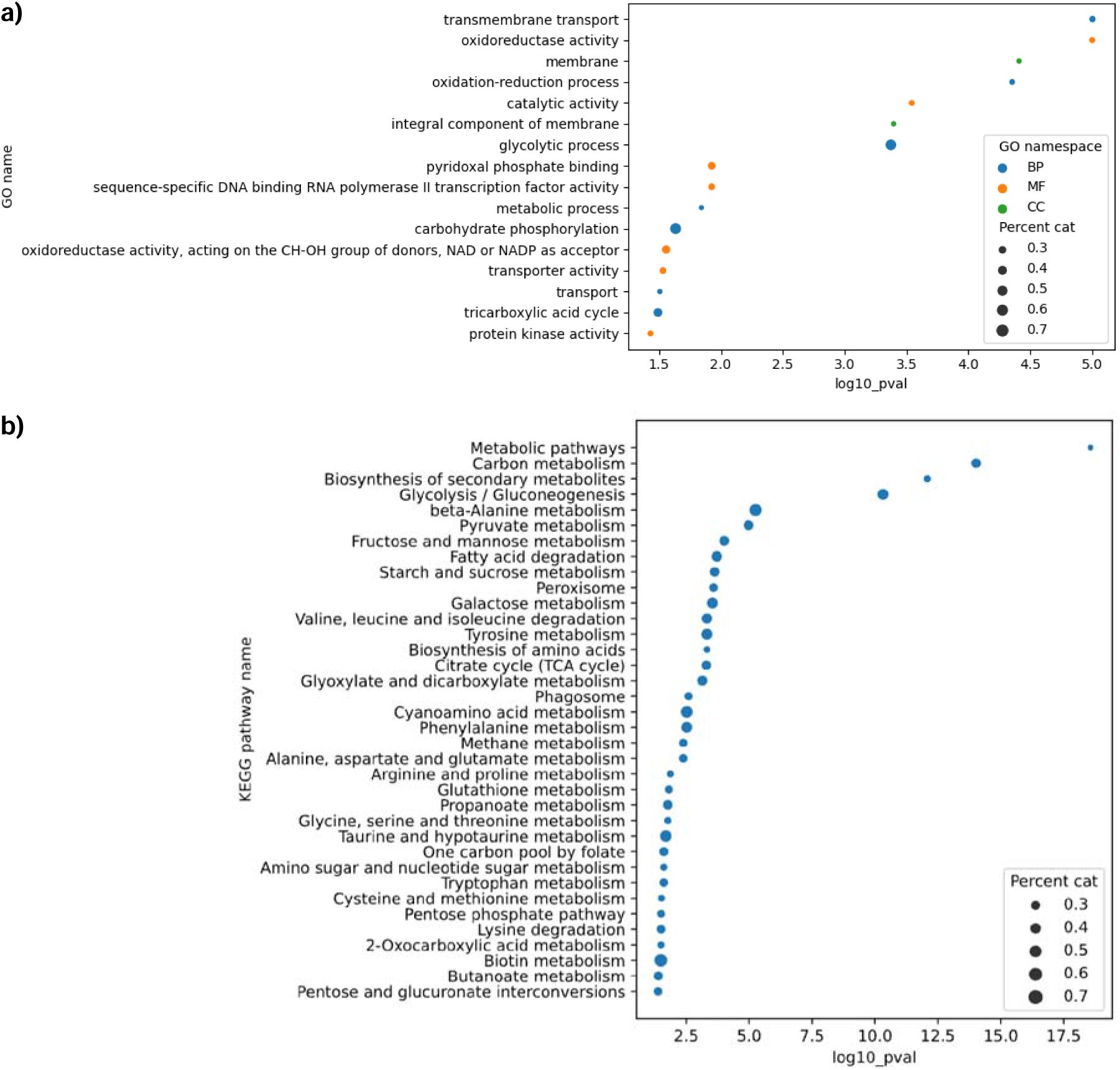
Significantly enriched categories identified through functional analysis when *D. hansenii* was exposed to 100 ppm of BaP. a) Gene ontology (GO) analysis and b) Kyoto Encyclopedia of Genes and Genomes (KEGG) pathway analysis. This analysis only includes overexpressed genes.

KEGG analysis revealed that 567 annotated ORFs were involved in key metabolic pathways, including amino acid biosynthesis, secondary metabolism, and energy production. Notably, genes linked to the glyoxylate shunt, methane metabolism, propanoate metabolism, and glutathione-mediated detoxification were strongly overrepresented (see Supplementary Figure 3 for a visual representation of all the KEEG maps that were identified as overrepresented under this specific condition; please refer to Supplementary Table 3 for a comprehensive list of these KEEG maps). These results suggest that BaP exposure does not only activate detoxification systems, but also triggers broader metabolic adjustments aimed at preserving homeostasis under stress.

To further elucidate the xenobiotic response mechanisms in *D. hansenii*, we conducted a focused transcriptomic analysis of 27 ORFs associated with canonical pathways involved in PAH degradation, glutathione (GSH) homeostasis, and oxidative stress defense (Figure 4). Gene selection was based on previous reports on PAH-degrading fungi and bacteria. Log_2_ fold changes (Log_2_FC) were calculated after 72 hours of incubation under BaP-only conditions, with glucose-grown cultures serving as the reference. As illustrated in Figure 4, the expression patterns of these genes closely paralleled the broader RNA-Seq trends, reinforcing their likely involvement in BaP detoxification and highlighting the role of GSH as a central adaptive mechanism.

**Figure 4.**
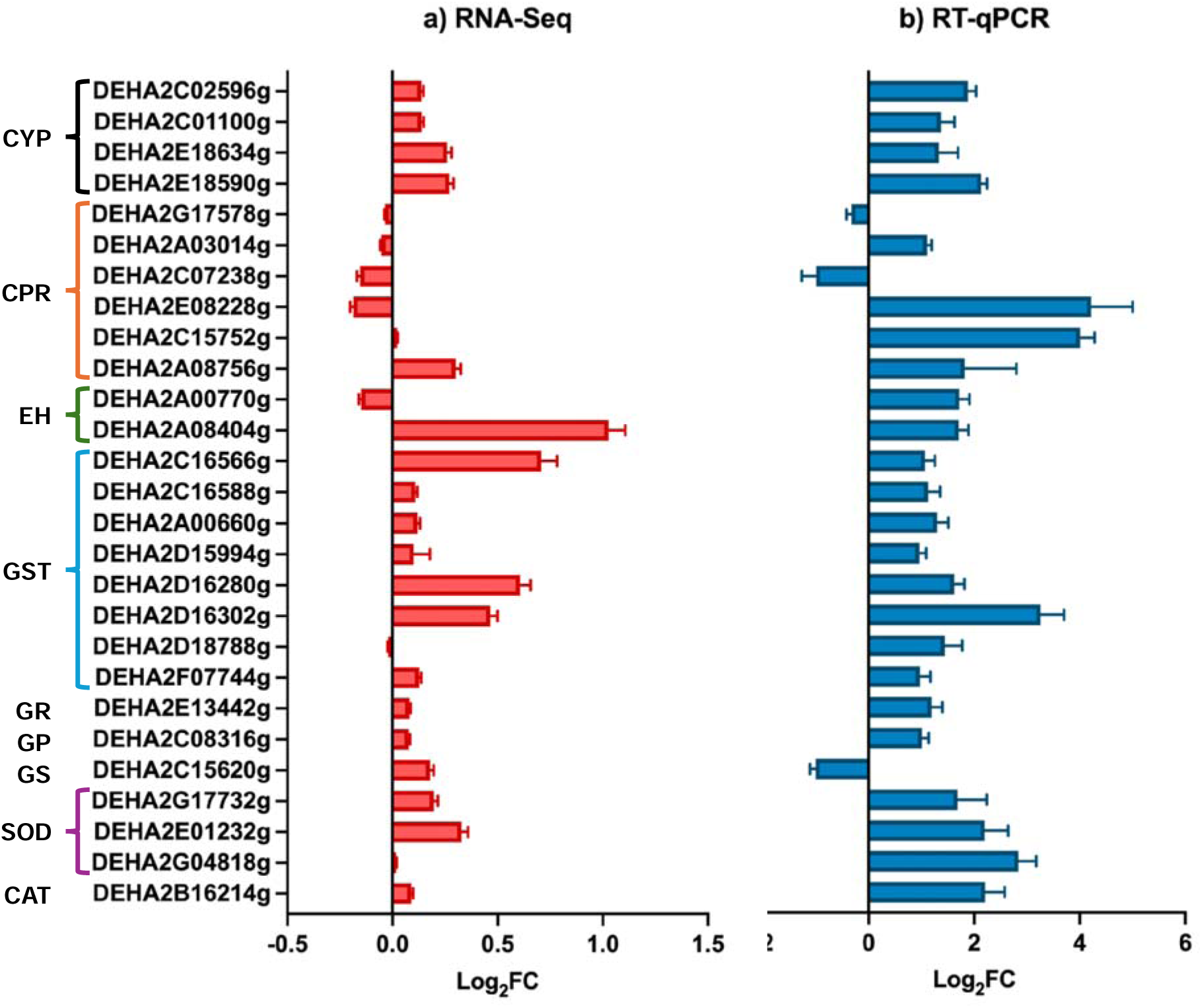
Relative expression of detoxification-related ORFs in *D. hansenii* exposed to 100 ppm of BaP. a) Differential expression analysis of ORFs obtained through RNA-Seq and b) Log_2_FC of relative expression in the presence of BaP using the *ACT1* housekeeping gene as a normalizer. Line colored brackets on the left axis, group ORFs belonging to a same metabolic process: CYP, cytochrome P450; CPR, cytochrome P450 reductase; EH, epoxide hydrolase; GST, glutathione S-transferase; GR, glutathione reductase; GP, glutathione peroxidase; GS, glutathione synthetase; SOD, superoxide dismutase; and CAT, catalase.

The strong induction of glutathione-related enzymes, alongside CYPs and EHs, underscores the importance of glutathione homeostasis and redox balance in *D. hansenii’s* response to BaP. RT-qPCR validation (Figure 4b) confirmed these expression trends, further supporting the robustness and consistency of the transcriptomic dataset.

The transcriptional changes observed in *D. hansenii* were specifically driven by BaP exposure rather than secondary to glucose deprivation, we conducted a second RNA-Seq experiment designed to control carbon source availability. In this assay, both control and treatment conditions included glucose (YNBG vs. YNBG + 100 ppm BaP), and samples were harvested at 72 hours to ensure consistency with the initial experiment. Library construction and sequencing were performed adhering to identical protocols. On average, sequencing yielded 47.2 million paired-end reads per sample (150 bp), with alignment rates approaching 100%, and 38% of reads mapping to annotated genes. These sequencing metrics closely matched those from the first dataset and were consistent with previous transcriptomic studies conducted under stress conditions (27).

PCA revealed a pronounced separation between control and BaP-treated groups, with biological replicates clustering tightly within each condition, demonstrating once again the reproducibility and robustness of the transcriptional response (Figure 5a).

**Figure 5.**
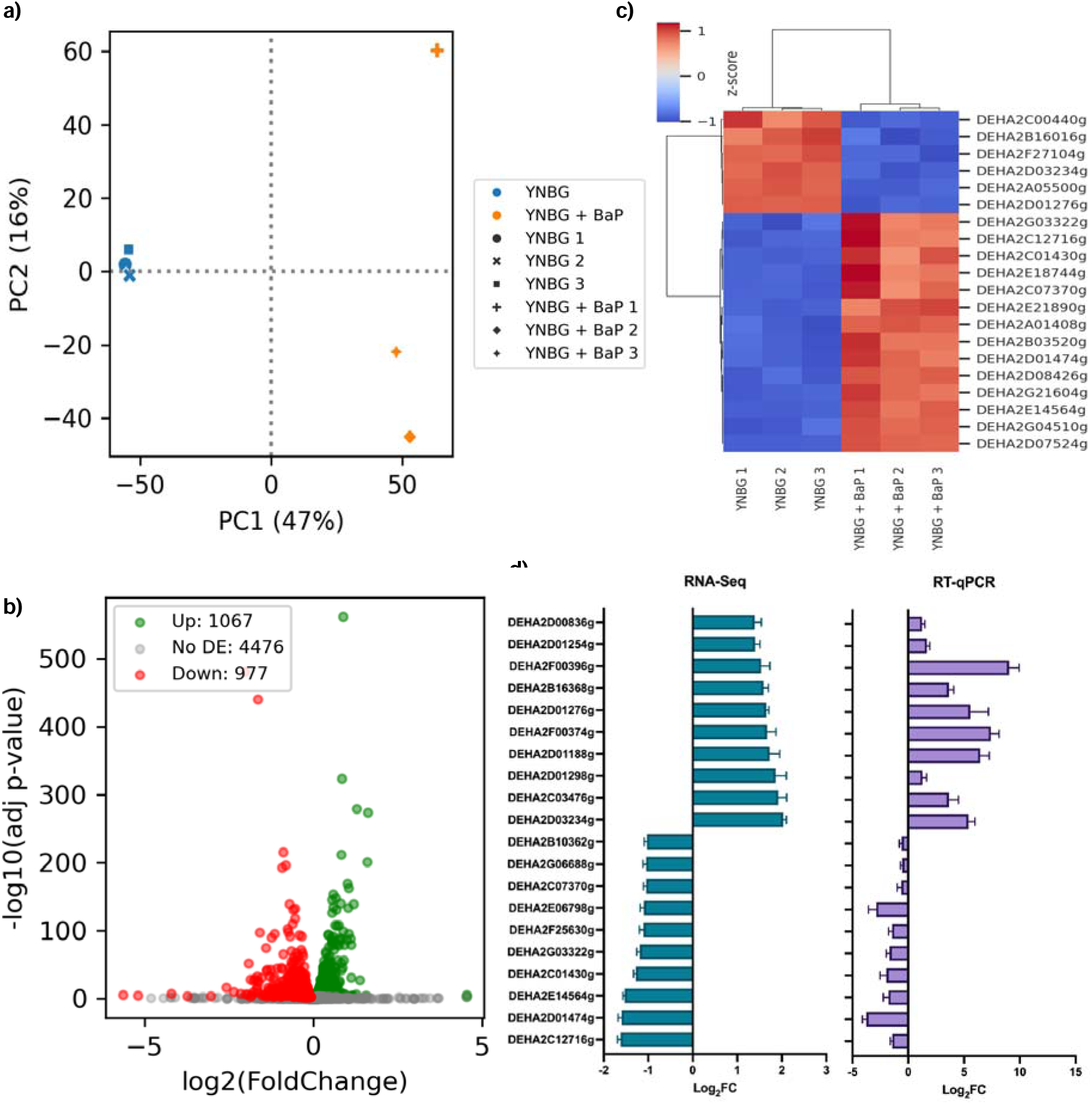
Transcriptomic analysis of *D. hansenii* exposed to glucose and 100 ppm of BaP. a) Principal component analysis YNBG control (YNBG 1, YNBG 2, and YNBG 3) and YNBG + BaP treatment (YNBG + BaP 1, YNBG + BaP 2, and YNBG + BaP 3). b) Volcano plot obtained from the differential expression analysis (adjusted *p*-value < 0.05). c) Top 20 most upregulated (red) and most downregulated (blue) genes based on RNA-Seq differential expression analysis (control condition: YNBG; experimental condition: YNBG + 100 ppm BaP). d) RT-qPCR validation of the 10 most overexpressed and the 10 most underexpressed genes identified through RNA-Seq differential expression analysis.

Differential gene expression analysis revealed 977 upregulated and 1067 downregulated genes under BaP treatment in glucose-supplemented conditions (Figure 5b). The most affected ORFs were predominantly linked to pathways involved in xenobiotic metabolism, redox homeostasis, membrane transport, and central carbon flux (Figure 5c). In contrast, downregulated genes showed significant enrichment in functional categories related to ion transport, protein folding, and broad-spectrum stress responses. Functional annotations were retrieved from InterPro (49). To validate these findings, we independently assessed the expression of the 10 most upregulated and 10 most downregulated ORFs via RT-qPCR using custom-designed oligonucleotides (Supplementary Table 1). As illustrated in Figure 5d, the resulting expression profiles closely reflected the RNA-Seq data, providing further support for the reproducibility of the BaP-induced transcriptional signature.

Following RT-qPCR validation, we performed functional annotation of the upregulated genes using GO and KEGG databases. GO term enrichment analysis (Figure 6a) highlighted overrepresentation in categories related to redox homeostasis, cellular stress responses, and various metabolic processes. In parallel, KEGG pathway mapping (Figure 6b) revealed that a substantial portion of these genes is involved in core metabolic routes, including the pentose phosphate pathway, the tricarboxylic acid cycle, the glyoxylate shunt, and lipid metabolism (see Supplementary Figure 4 for a visual representation of all the KEEG maps that were identified as overrepresented under this specific condition; please refer to Supplementary Table 4 for a comprehensive list of these KEEG maps). Together, these results suggest that *D. hansenii* orchestrates a coordinated reprogramming of its metabolism when challenged with BaP, redirecting both energy flow and redox capacity toward detoxification and stress adaptation.

**Figure 6.**
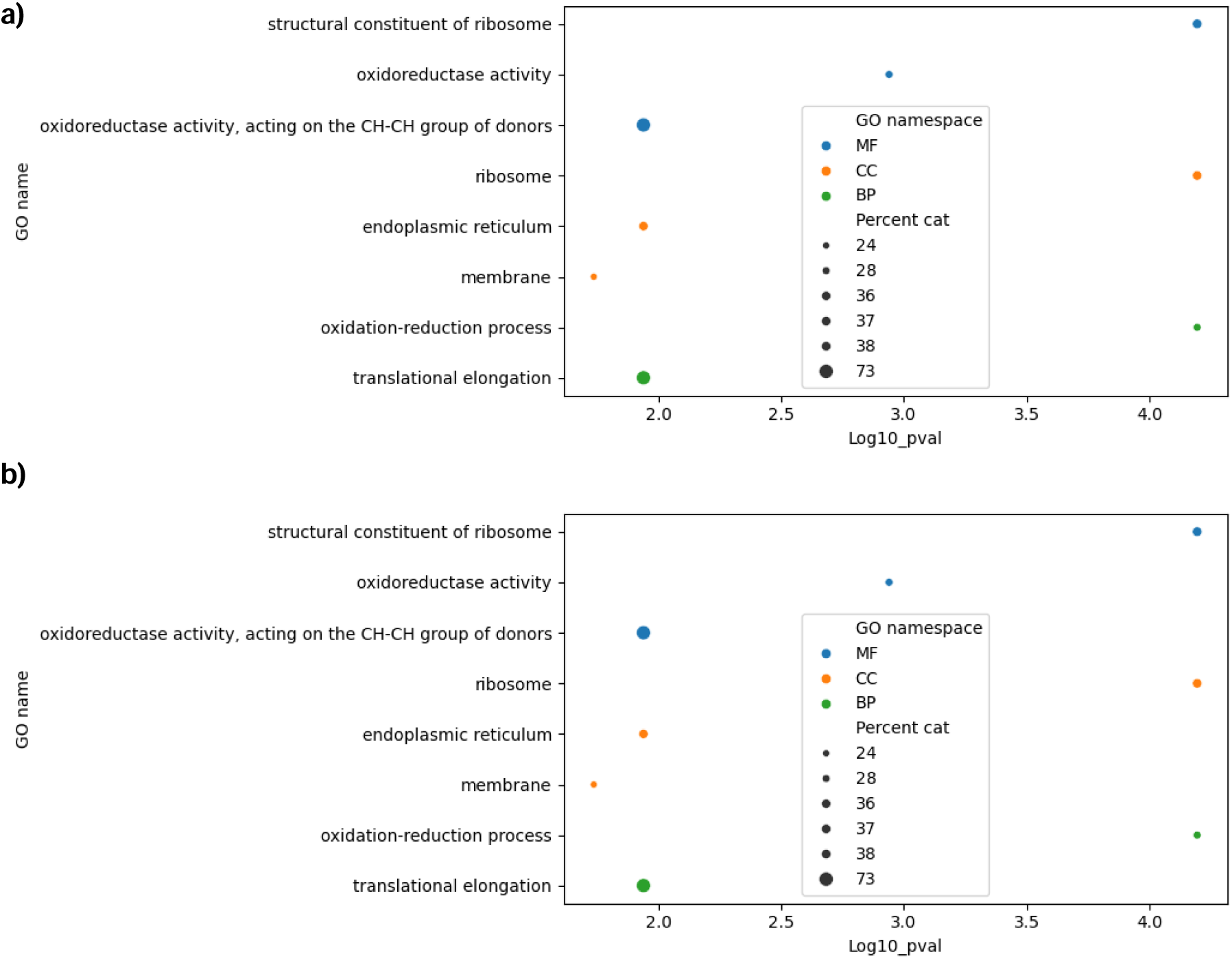
Significantly enriched categories identified through functional analysis. a) Gene ontology (GO) analysis and b) Kyoto Encyclopedia of Genes and Genomes (KEGG) pathway analysis. This analysis only includes overexpressed genes.

However, the changes were more pronounced in the absence of glucose, which is somewhat expected given that carbohydrate catabolism is favored under glucose-rich conditions. In contrast, when carbohydrates such as glucose are absent, anabolic processes tend to predominate, thereby enhancing the activity of the aforementioned metabolic pathways.

To determine whether glucose modulates the detoxification response, we re-examined the expression of five key ORFs previously identified as BaP-responsive (Figure 4), now under co-metabolic conditions (YNBG + BaP vs. YNBG). After 72 hours of exposure, Log_2_FC revealed expression patterns remarkably similar to those observed under BaP-only conditions (Figure 7), suggesting that glucose does not suppress, but may instead support, the activation of xenobiotic metabolism and glutathione homeostasis pathways. RT-qPCR validation of these genes confirmed the trends seen in the transcriptomic data, with minor discrepancies likely attributable to technical variation between platforms. These findings reinforce the central role of CYPs, GSTs, and antioxidant systems in *D. hansenii*’s response to BaP, highlighting a robust and conserved detoxification strategy regardless of carbon source availability (50).

**Figure 7.**
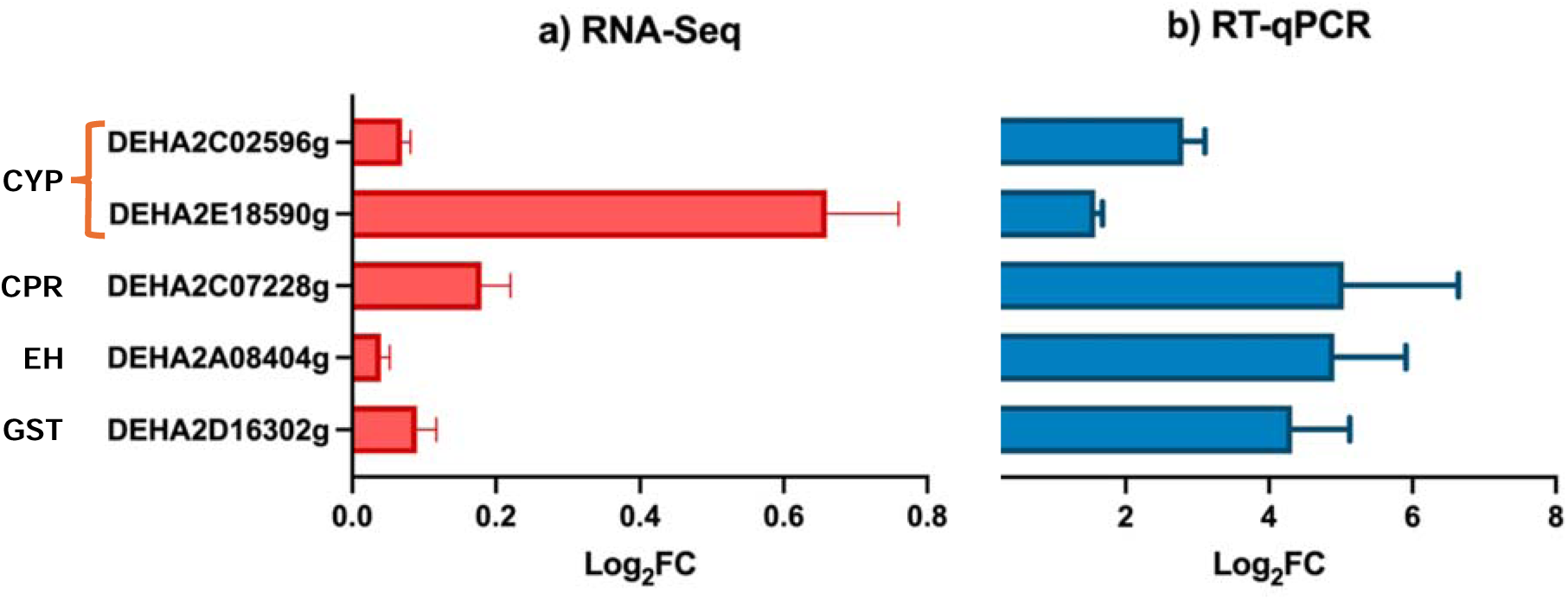
Relative expression of detoxification-related ORFs in *D. hansenii* exposed to BaP. a) Differential expression analysis of ORFs obtained through RNA-Seq and b) Log_2_FC of relative expression in the presence of glucose+ BaP using the *ACT1* housekeeping gene as a normalizer. Line colored brackets on the left axis, group ORFs belonging to a same metabolic process: CYP, cytochrome P450; CPR, cytochrome P450 reductase; EH, epoxide hydrolase; GST, glutathione S-transferase.

Furthermore, exposure to BaP + glucose led to the strong induction of four CYP genes and their corresponding reductases, enzymes involved in the initial oxidation of BaP into reactive intermediates such as epoxides. These transcriptional changes coincided with the upregulation of EHs and GSTs, which mediate the conversion and conjugation of these intermediates into more stable, less toxic products.

Simultaneously, the yeast activated genes coding for antioxidant enzymes, including superoxide dismutase (SOD), catalase (CAT), and glutathione peroxidase (GP) (Figure 4), that help neutralization of reactive oxygen species generated during BaP metabolism. Together, these expression patterns reflect the deployment of a coordinated cellular program that integrates enzymatic detoxification with redox defense mechanisms in *D. hansenii* exposed to BaP.

In summary, our physiological and transcriptomic analyses revealed that *D. hansenii* maintains growth and efficiently degrades BaP under both nutrient-limited and co-metabolic conditions. The yeast activates a coordinated detoxification program encompassing xenobiotic metabolism, antioxidant defenses, membrane transport, and metabolic reprogramming. These responses were consistent across two independent RNA-Seq experiments, highlighting the robustness and reproducibility of *D. hansenii*’s adaptation to BaP exposure (Supplementary Table 2). Together, these findings provide a comprehensive molecular framework for understanding the mechanisms underpinning *D. hansenii*’s survival mechanisms and set the foundation for the discussion of its potential as an extremophilic bioremediation agent.

## 4. Discussion

This study provides an integrated physiological and transcriptomic perspective on *D. hansenii*’s response to BaP, highlighting its remarkable resilience and metabolic flexibility. The yeast not only tolerates BaP but maintains growth and achieves substantial degradation under both nutrient-limited and co-metabolic conditions (Figure 1a), in agreement with previous findings (21, 23). Notably, its reported capacity to survive up to 500 ppm BaP, alongside its ability to metabolize other hydrocarbons such as n-dodecane and thrive in high-salinity, cold, or metal-stressed environments, further supports its promise as a bioremediation agent in extreme ecological niches (26, 27, 51).

Growth and biomass assays (Figures 1b and 1c) revealed that *D. hansenii* grows more slowly and accumulates less biomass when BaP is the sole carbon source, likely due to the energetic cost associated with detoxification processes. This oligotrophic adaptation, also described in *Candida albicans*, *Saccharomyces cerevisiae*, and *R. mucilaginosa*, supports survival under nutrient stress and aligns with the broader concept of metabolic trade-offs under toxic exposure (23, 28, 52–54).

Under co-metabolic conditions, glucose supplementation enhanced both biomass production and BaP degradation efficiency, increasing from 65.0% in minimal medium to 73.1% with glucose (Figure 1d). This pattern mirrors observations in other microbes, such as *Armillaria* sp., *Acinetobacter johnsonii*, and *Pichia anomala*, where glucose acts as a co-substrate that fuels detoxification enzymes (19, 20, 55). Interestingly, glucose consumption decreased in BaP-exposed cultures (Figure 1e), suggesting either a redirection of metabolic activity toward BaP degradation or a chemical stress-induced slowdown of glycolytic flux (56).

The metabolic flexibility of *D. hansenii* is further evidenced by its ability to utilize hydrophobic substrates in hydrocarbon-rich environments, a trait it shares with other extremotolerant organisms (21, 23, 51). Such metabolic rerouting has also been observed in bacteria, where exposure to xenobiotics redirects carbon and energy fluxes toward detoxification, often at the expense of growth (57–59).

In nutrient-limited media (YNB + BaP), *D. hansenii* appears to favor survival over proliferation by activating detoxification pathways and stress maintenance programs rather than investing in biomass accumulation. This strategy reflects a broader adaptive principle observed in multiple fungi and yeasts exposed to PAHs (21, 23, 60).

This work offers the first transcriptomic insight into the *D. hansenii* response to BaP, revealing a tightly regulated network of detoxification, redox control, and metabolic adaptation (see Supplementary Table 2). Similar transcriptional or proteomic responses have been described in *Aspergillus* spp., *R. mucilaginosa*, *S. cerevisiae*, *Dentipellis* sp., and in the marine alga *Ulva lactuca* under xenobiotic stress (9, 28, 29, 50, 53, 61). These parallels suggest a conserved eukaryotic strategy to cope with aromatic hydrocarbon toxicity.

Nutrient availability, particularly glucose, clearly modulated transcriptional responses. Principal component analysis (Figure 2a and 5a) revealed that nutrient-limited conditions produced a more uniform expression pattern, while co-metabolic conditions triggered a broader, more dynamic response (62). This distinction underscores the dual role of glucose as both an energy source and a modulator of detoxification under chemical stress (63, 64).

Functional enrichment (Figures 3 and 6) showed significant overrepresentation of genes involved in xenobiotic metabolism, redox homeostasis, the glyoxylate cycle, carbohydrate processing, and membrane transport. These metabolic modules appear to form the molecular foundation of *D. hansenii’s* adaptive response to BaP, facilitating detoxification and stress resistance (Supplementary Table 2).

Across conditions, *D. hansenii* consistently activated7 key pathways related to xenobiotic degradation, antioxidant defense, and carbon rerouting—particularly the glyoxylate cycle, which supports energy conservation under stress. This metabolic shift resembles responses observed in *S. cerevisiae* exposed to acetic acid (65, 66), *Candida glabrata* within macrophages (67, 68), and various bacteria under oxidative or chlorine stress (69, 70).

Phase I detoxification was especially robust, as evidenced by the strong upregulation of cytochrome P450 monooxygenases (especially *DhDIT2*, DEHA2C02596g), and their reductases (CPRs), which mediate the initial oxidation of BaP. These enzymes convert BaP into reactive epoxides, which are subsequently transformed by EHs into less harmful diols. This sequential enzymatic strategy mirrors fungal detoxification pathways reported in *Aspergillus* spp. and *R. mucilaginosa* (23, 28, 29, 61, 71).

The Phase II response featured increased expression of GSTs, glutathione reductase (GR), glutathione peroxidase (GP), and glutathione synthetase (GS), enabling the conjugation of electrophilic intermediates and the recycling of reduced GSH (72). This system was supported by the upregulation of antioxidant enzymes such as SOD and CAT, which collectively manage ROS levels generated during BaP metabolism (73).

Comparable glutathione-antioxidant systems have been described in *C. albicans* and *Penicillium chrysogenum*, where detoxification is linked to NADPH production via the pentose phosphate pathway (74, 75). Although the GABA shunt was not significantly upregulated in *D. hansenii*, the transcriptomic data suggest a compensatory reliance on the pentose phosphate and glyoxylate cyclesas primary sources of NADPH and biosynthetic precursors under stress (73, 76, 77).

In addition to detoxification, *D. hansenii* undergoes substantial metabolic reprogramming under BaP stress. Upregulated genes involved in the glyoxylate cycle, β-oxidation, and the TCA cycle indicate a shift in carbon flux to preserve ATP generation and redox homeostasis. Prior studies have shown that *D. hansenii* also activates the glyoxylate cycle under salt stress, suggesting that this pathway may represent a core survival mechanism across different environmental insults (78), similar to what occurs in the presence of BaP. These patterns align with adaptations observed in *Aspergillus* spp. under glucose-rich conditions and in *R. mucilaginosa* exposed to PAHs, where energetic metabolism shifts support stress endurance (28, 61).

Structurally, *D. hansenii* exhibits upregulation of ATP-binding cassette (ABC) and Major Facilitator Superfamily (MFS) transporters, alongside genes involved in lipid remodeling, suggesting not only active export of BaP-derived metabolites but also a concerted adaptation to maintain membrane stability under xenobiotic stress. Similar strategies have been reported in *Fusarium solani*, *Yarrowia lipolytica*, and *Trichosporon* spp., where PAHs interact with membrane bilayers, prompting changes in lipid composition and transporter activity to facilitate pollutant efflux and limit membrane permeability (79–82). These alterations are essential for survival in environments rich in hydrophobic toxics like BaP, which tend to accumulate in cellular membranes and impair their function (7).

Although *AOX1* expression was undetectable, transcriptional signatures indicative of mitochondrial metabolic changes were evident. This may reflect the activation of alternative energetics strategies, such as the activation of the glyoxylate pathway. In *Candida parapsilosis*, for instance, exposure to hydroxyaromatic compounds leads to the induction of mitochondrial carriers that connect xenobiotic degradation to central carbon metabolism, ensuring energy production under stress (83). Moreover, it has been shown that PAHs can disrupt mitochondrial membrane potential and function, leading to compensatory adjustments in respiratory chain components, antioxidant defenses, and metabolic fluxes, as described in yeast and mammalian systems (84, 85). These parallels suggest that *D. hansenii* may adopt a comparable metabolic reprogramming to mitigate the deleterious effects of BaP at the organelle level.

The coordinated upregulation of CYP monooxygenases, CPRs, EHs, GSTs, and antioxidant enzymes highlights an integrated detoxification system in *D. hansenii*, combining phase I oxidation, phase II conjugation, and redox balance. This orchestrated response enables the yeast to process BaP-derived intermediates efficiently while minimizing oxidative damage (10, 23). Similar integrative detoxification strategies have been observed in *Ulva lactuca*, *A. sydowii*, and *R. mucilaginosa* under PAH exposure, supporting the notion of a conserved eukaryotic framework for PAHs and other xenobiotic defense (11, 28, 29, 50, 86, 87).

What truly distinguishes *D. hansenii* is its ability to thrive where most bioremediators fail. Salinity, cold, and metal-rich environments, harsh conditions typical of polluted sites, don’t significantly hinder its growth or function. This extremotolerance, well-documented in previous studies (26, 27, 51), adds a critical layer of ecological relevance to its already impressive metabolic versatility. Much like *Candida parapsilosis*, which adjusts its mitochondrial dynamics under xenobiotic stress (83, 84), *D. hansenii* appears to modulate both its organelles and membranes to remain viable and metabolically active in toxic environments.

Our findings converge in a five-tiered model (Figure 8) that captures the structured and adaptive response of *D. hansenii* to BaP exposure. This model integrates xenobiotic sensing, antioxidant defense, conjugation and export of metabolites, and metabolic reprogramming into a cohesive framework that reflects both conserved eukaryotic strategies and the unique extremotolerance of this yeast. Crucially, this response remains effective even under nutrient-limited conditions, underscoring *D. hansenii*’s potential for survival and biotransformation in harsh, contaminated environments.

1. Xenobiotic activation via CYPs and CPRs, leading to the formation of reactive epoxides.
2. ROS generation as a consequence of BaP oxidation and detoxification of reactive intermediates through conjugation with GSH by GSTs, alongside the upregulation of genes involved in GSH synthesis and recycling.
3. Elimination of conjugates via ABC and MFS transporters or internalization in the vacuole.
4. Maintenance of redox and energetic balance through the activation of the pentose phosphate pathway, glyoxylate cycle, and other metabolic processes.
5. Membrane remodeling.

**Figure 8.**
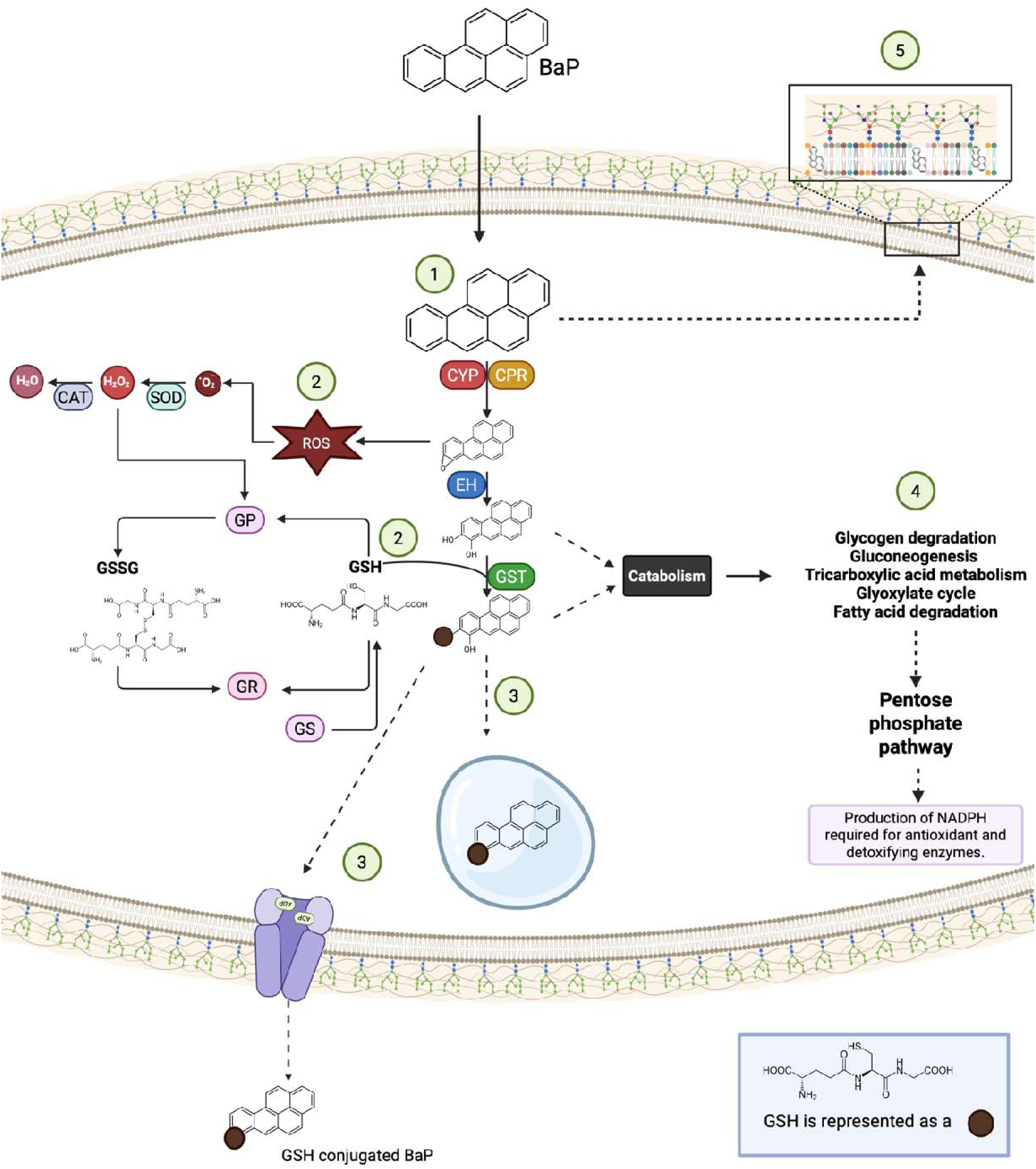
Five-tier model summarizing the adaptive response *of D. hansenii* to BaP exposure. After internalization, (1) BaP is activated via cytochrome P450 monooxygenases (CYP), NADPH-cytochrome P450 reductases (CPR), and epoxide hydrolases (EH); (2) antioxidant defense through glutathione-related enzymes (glutathione S-transferases (GST), glutathione reductase (GR); glutathione peroxidase (GP), glutathione synthetase (GS)) and ROS-scavenging enzymes (superoxide dismutase (SOD) and catalase (CAT)); (3) metabolite conjugation and export by ATP-binding cassette (ABC) and Major Facilitator Superfamily (MFS) or vacuolar internalization; (4) metabolic reprogramming involving the pentose phosphate pathway, glyoxylate cycle, β-oxidation, and the tricarboxylic acid cycle to maintain redox balance; and (5) membrane remodeling. This figure was constructed using data from KEGG pathways that are significantly overrepresented in the presence of BaP (see Supplementary Figures 3 and 4).

## 5. Conclusions

This study confirms that *D. hansenii* can both grow and degrade BaP under nutrient-limited and glucose-supplemented conditions, demonstrating its exceptional metabolic adaptability in stressful environments. Although BaP alone supports growth and degradation, co-metabolism with glucose significantly improves biomass accumulation and degradation efficiency.

Transcriptomic analyses revealed that both conditions triggered canonical detoxification responses, including CYP-mediated oxidation, GSH-based conjugation, antioxidant enzyme activation, and membrane remodeling. The co-metabolic condition (YNBG + BaP) further promoted energy-producing pathways such as the pentose phosphate pathway, which is essential for NADPH generation and efficient detoxification. While the GABA shunt was anticipated to play a role in redox balance, its transcriptional regulation was not evident, suggesting the activation of alternative compensatory mechanisms.

Overall, our findings position *D. hansenii* as a uniquely equipped extremophilic yeast, capable of metabolizing BaP through a tightly coordinated transcriptional program that integrates detoxification, antioxidant defense, membrane remodeling, and metabolic plasticity. This multifaceted response supports its candidacy as a robust platform for bioremediation, particularly in environments where conventional microbial systems may fail.

## Supporting information

All suplemental

## 6. Acknowledgements

The authors gratefully acknowledge Dr. Diego González-Halphen and Dr. James González for their valuable scientific guidance. We also thank Dr. Laura Ongay-Larios, MSc. Minerva Mora, BSc. Guadalupe Codiz-Huerta, Dr. Carlos Alberto Peralta-Álvarez, Dr. Martha Lucinda Contreras-Zentella, Dr. Carlos Campero-Basaldua and Eduardo Cardona Guerrero for their technical assistance. The support of Dra. Claudia Segal-Kischinevzky, Dra. Alicia González Manjarrez, Dr. Roberto Arreguín-Espinosa, and Dr. Daniel Genaro Rosas with the performance of some the experiments in this work is also gratefully acknowledged.

This manuscript was written by the authors. Generative AI tools (ChatGPT-4 and DeepL Write) were only used to refine language and clarity, with all scientific content authored and reviewed by the authors. Additionally, Mendeley was employed for the management of bibliographic references, and BioRender.com was employed to create Figure 8.

## 7. Author Contributions

Conceptualization: F.P.-G., A.P.; Data curation: F.P.-G., A.C.P.-H.; Formal analysis: F.P.-G., A.C.P.-H.; Funding acquisition: A.P.; Investigation: F.P.-G., M.A.-V., M.C., N.S.S.; Methodology: F.P.-G.; Project administration: A.P.; Resources: A.P.; Software: A.C.P.-H.; Supervision: A.P.; Validation: F.P.-G., M.A.-V., M.C., N.S.S., A.P.; Visualization: F.P.-G., A.C.P.-H.; Writing – original draft: F.P.-G., A.P.; Writing – review & editing: F.P.-G., A.C.P.-H., M.A.-V., M.C., N.S.S., A.P. All authors contributed to revising the manuscript.

## 8. Fundings

This research was supported by grants IN204321 and IN217924 from the Dirección General de Asuntos del Personal Académico (DGAPA), Universidad Nacional Autónoma de México (UNAM).

Francisco Padilla-Garfias is a doctoral student in the Programa de Maestría y Doctorado en Ciencias Bioquímicas at UNAM and receives a scholarship (CVU 904691) from the Secretaría de Ciencia, Humanidades, Tecnología e Innovación (SECIHTI). The authors thank DGAPA-UNAM for the fellowship provided to Minerva Georgina Araiza-Villanueva.

## 9. Ethics Statement

All experimental procedures involving *Debaryomyces hansenii* were carried out in accordance with institutional biosafety guidelines of the Instituto de Fisiología Celular, UNAM. The yeast strain used in this study is non-pathogenic and does not require special ethical approval under current national or international regulations.

## 10. Conflicts of Interest

The authors declare no conflicts of interest.

## Notes

### Competing Interest Statement

The authors have declared no competing interest.

