## Supplementary material for "Transcriptomic profiling of *Debaryomyces hansenii* reveals detoxification and stress responses to benzo(a)pyrene exposure": All suplemental: Supplementary_Table_1.docx

**Supplementary Table 1. Oligonucleotides used in qPCRs**

| **ORF** | **Oligonucleotide** | **Secuence** |
| --- | --- | --- |
| **DEHA2D05412g**  ***ACT1*** | Forward | CCCAGAAGAACACCCAGTTT |
|  | Reverse | CGGCTTGGATAGAAACGTAGAA |
| **DEHA2G00132g** | Forward | TGATCGTATAGGAAGAAAGCACA |
|  | Reverse | TTCCAGTTGCACCAGAACCA |
| **DEHA2E01166g** | Forward | TTAACTGGAGCTGCGCTCAT |
|  | Reverse | ATGCAATAACAGCCTTGGCG |
| **DEHA2E00176g** | Forward | GCCCCAGAATTTTACGGCAG |
|  | Reverse | CCGCGCTGGGAAGATACATA |
| **DEHA2A06314g** | Forward | TGTCTATCCCGACACCTCCA |
|  | Reverse | TGGTGTTTCTCAGCTGCCAC |
| **DEHA2C16324g** | Forward | ACCTGCTCAAAGCCAATCCA |
|  | Reverse | CTCAATAAGCAACGCGGCAA |
| **DEHA2B05390g** | Forward | ACAGACCCCTAGACCGTTGA |
|  | Reverse | GGTACAACCGCCAGTCTTGA |
| **DEHA2F27390g** | Forward | AGCTGATCCACTTAATGATCTTTTT |
|  | Reverse | TGCATATCTGAATCCGACTCGT |
| **DEHA2C05918g** | Forward | CGCATGAAGAGAATGGGTTGT |
|  | Reverse | GCATCACCTGCTTCTCGGA |
| **DEHA2B15928g** | Forward | ACCCCATGTCCATACTGGGA |
|  | Reverse | GGTAGCTAAGCCAGCAGCAT |
| **DEHA2F17996g** | Forward | GAGCCACCTGTAGCAGAAGT |
|  | Reverse | GCTGCTAAACCCAAAGGTGC |
| **DEHA2G24200g** | Forward | ATGCCAATTTCACCGGATACGA |
|  | Reverse | GACATGTTCTTGGTGACACGC |
| **DEHA2F27016g** | Forward | ACATGGCCGCTCACCATATC |
|  | Reverse | AAAGCACCAAAAGCGCATCC |
| **DEHA2D00330g** | Forward | TGATTGCGGAGGCTACGTTC |
|  | Reverse | GCCGACCCTACCTAAAAACC |
| **DEHA2D01232g** | Forward | GGCAAGGAAGCTCATAGCGA |
|  | Reverse | CTGGATCATCAGGCCCGTC |
| **DEHA2G20746g** | Forward | TGTCTGGCTGCTTGATGGAG |
|  | Reverse | CCAGTCAATCGTTCCTGGCT |
| **DEHA2F22660g** | Forward | TTCGTTGCTGGAGACGTCAA |
|  | Reverse | GCTCCCATTGTTCGGTCGTA |
| **DEHA2D01364g** | Forward | ACGTACTATTCGGATTCGCCA |
|  | Reverse | CAGACATGTGGACCCGRGAG |
| **DEHA2B16478g** | Forward | CTCCAAGCATCTGTGGAGCA |
|  | Reverse | TGTGCGTCGGAAACAGATGA |
| **DEHA2B00286g** | Forward | TGAAAGGTGATTGGGGGTCAG |
|  | Reverse | ACCATGCCCGCTATTTGAC |
| **DEHA2E01188g** | Forward | TGGTCGTCTGGTTATGCTGC |
|  | Reverse | AACCAGTGGCAGGATGTGTT |
| **DEHA2C01100g** | Forward | CGATGCAGCAACGGAGTTTT |
|  | Reverse | TACAAATCCTGCGCCAAAGC |
| **DEHA2E18634g** | Forward | TCGGAGAGGGAGAAGATTCCA |
|  | Reverse | TTGGCGATTTTCCATCGGGA |
| **DEHA2E18590g** | Forward | TTGGTGCGAAACCAATGCAG |
|  | Reverse | CATCCCGCGTAACTACAGCA |
| **DEHA2G17732g** | Forward | CGGTGTCGTTAACTTCGAACA |
|  | Reverse | TGAACGGGTTAAAGTGAGGT |
| **DEHA2E01232g** | Forward | GTGGAAAACATCCAGGATCGG |
|  | Reverse | AGGTTCCCACCGTTCTGTTT |
| **DEHA2G04818g** | Forward | CGTCGGACTCTGAATTGGCA |
|  | Reverse | TGGTCCCGTAACAGTGTCTTG |
| **DEHA2B16214g** | Forward | CATAYGGTGCCCAAACGGC |
|  | Reverse | ACCACCACCTTTTGCGTGAA |
| **DEHA2E13442g** | Forward | ACCTTAACCGGAGACCAAGA |
|  | Reverse | TAGCATATCCTCCTGTGGCA |
| **DEHA2C08316g** | Forward | GAGAGGGGATTTGTTGTGCT |
|  | Reverse | TCCGTTGCTCGTTCTTCAAA |
| **DEHA2C15620g** | Forward | ACTCACAGTGCTACACATGC |
|  | Reverse | AGTCATCGTCAAAAACGGCT |
| **DEHA2C02596g** | Forward | GCGTTGGGTACAAGACATGC |
|  | Reverse | CTGTTCACGTGCAAACTGGG |
| **DEHA2G17578g** | Forward | CCGATTACGTATGGTGCCGT |
|  | Reverse | ATCTCCGTGGCAAACAACTCT |
| **DEHA2A03014g** | Forward | AAATCGACGGAAAGGACGAG |
|  | Reverse | AGCATGGCAAATCTCTTGGA |
| **DEHA2C07238g** | Forward | ACTCCAATCTCCACCGATGA |
|  | Reverse | CGATTCAACCATACCTGGGG |
| **DEHA2E08228g** | Forward | ACAAGGCTCCTGAAAACTGG |
|  | Reverse | GGACCACAGACGAAAACCTT |
| **DEHA2C15752g** | Forward | GGGTGGTCATGGTGTTTACC |
|  | Reverse | TCCCATTGCTTGCCATACTT |
| **DEHA2A08756g** | Forward | GGTGCACTTGGTATGATGGT |
|  | Reverse | CTGGAACAAGTCTCCAGCAA |
| **DEHA2D15994g** | Forward | CGTTCTCTCCCACCAAAGGC |
|  | Reverse | TGGATTCTGGGTCAGCAGAGT |
| **DEHA2D16280g** | Forward | AGGCCCTCGACTTTTTGGAA |
|  | Reverse | CCGCCAGTCATAGCCTTTGT |
| **DEHA2D16302g** | Forward | AGAAAGGGAGGGGACAGGTT |
|  | Reverse | CACCATGCAGCAAGATGTGG |
| **DEHA2D18788g** | Forward | TTGTTAAGTTCGCGCCTGAG |
|  | Reverse | ACTTGGCCCACGAATAGACG |
| **DEHA2F07744g** | Forward | GCGGCCGCTTATAGTGCTG |
|  | Reverse | GCCACAGTAGCTCTATCACCAACT |
| **DEHA2D03234g** | Forward | TTTGCTTTGCATGGTGTTGG |
|  | Reverse | CACATGAGCCTGCAATTTGG |
| **DEHA2C03476g** | Forward | AAAGTGGGTTGGTCGTCATC |
|  | Reverse | AATTCGAAACCTTCGTCCCC |
| **DEHA2D01298g** | Forward | AGTCATGGTAAGCGCATTCA |
|  | Reverse | CCATATTGCGACAGCTCCAT |
| **DEHA2D01188g** | Forward | TTGGACAACTAACACCACCG |
|  | Reverse | GCACCCAATTCAATCGCAAA |
| **DEHA2F00374g** | Forward | GTGGACATTCAAGGGAGCAT |
|  | Reverse | AGACAACTCTTCCATTCGCC |
| **DEHA2D01276g** | Forward | TCGGTCACAAGAGAAAGCAG |
|  | Reverse | TGCCAACACTTTTGCTGATG |
| **DEHA2B16368g** | Forward | TTGAGGATATTGCTTGCGCT |
|  | Reverse | TACCAATCCAGACCCACACT |
| **DEHA2F00396g** | Forward | AAACGTTTTGGTTGCTACGG |
|  | Reverse | CGGATCCCCTTTCATTCTGG |
| **DEHA2D01254g** | Forward | TATTTGAGACACCCGGAGGA |
|  | Reverse | AGGACCATTTTTGGCACTTCT |
| **DEHA2D00836g** | Forward | TAGGCTATTTGGCTGTGGGA |
|  | Reverse | ACACCCTGATATTAGTCACTCCT |
| **DEHA2C12716g** | Forward | ATCAGGCGCTCCATTTAGTG |
|  | Reverse | GGAGCTTGTGGACCTTCTTC |
| **DEHA2D01474g** | Forward | GATAGTGCCGACGCTAACAA |
|  | Reverse | GAAGCGGCAGATAATGCAAC |
| **DEHA2E14564g** | Forward | GTGGAATTCCTTCCCCATCC |
|  | Reverse | GACTTTTCTGGGTCTGCCAT |
| **DEHA2C01430g** | Forward | CAACATAGCTGCTGCGAAAG |
|  | Reverse | AAGAACAACTCTTGCGACGA |
| **DEHA2G03322g** | Forward | GTCCCCATCTCCAGACCTTA |
|  | Reverse | GTCTTGCCATAGCAGGAACA |
| **DEHA2F25630g** | Forward | CGGGAGGAAGTGGAAAGATG |
|  | Reverse | AGTGGCAATACTGGCTGATG |
| **DEHA2E06798g** | Forward | TCCTCCGGAAGTTCATTGGA |
|  | Reverse | ACGTCATGGATGAGGTACGA |
| **DEHA2C07370g** | Forward | AATGATCGACCAAAAGGCCA |
|  | Reverse | ATTGAAAGCAGCAGCCAAAG |
| **DEHA2G06688g** | Forward | TTCCGATGGTCAAACCACAG |
|  | Reverse | TCATCATCGTCATCGGTTGC |
| **DEHA2B10362g** | Forward | AGAACTGGAAGTTGTGCTGG |
|  | Reverse | CGAGTGCACCTCTTAATGCT |
