## Supplementary material for "Transcriptomic profiling of *Debaryomyces hansenii* reveals detoxification and stress responses to benzo(a)pyrene exposure": All suplemental: Supplementary_Table_2.docx

**Supplementary Table 2. ORFs of interest**

| **ORF** | **Function of the ORF** |
| --- | --- |
| **Detoxifying ORFs** | |
| DEHA2A04554g | Xenobiotic transmembrane transporter activity. |
| DEHA2A10362g | The dityrosine exporter, Dtr1 (required for formation of the outer layer of the cell wall. |
| DEHA2B06644g | The Hol1 MFS transporter. |
| DEHA2B14740g | Cycloheximide: H^+^ antiporter. |
| DEHA2C00858g | Xenobiotic transmembrane transporter activity. Uncharacterized membrane protein, predicted efflux pump. |
| DEHA2C03718g | The main boron exporter in yeast, Atr1. |
| DEHA2C05390g | Siderophore-iron (ferrioxamine):H^+^ symporter, Sit1 (Arn3) (in vesicles). |
| DEHA2C10274g | Glucose-6-phosphate dehydrogenase activity. |
| DEHA2C16654g | Antioxidant activity. Alkyl hydroperoxide reductase, thiol specific antioxidant and related enzymes. |
| DEHA2D07832g | Monoatomic anion transport. |
| DEHA2D09768g | Xenobiotic transmembrane transport, uncharacterized membrane protein, predicted efflux pump. |
| DEHA2D11506g | Response to stress, the 7-amino cholesterol resistance protein, Rta1p. |
| DEHA2D16896g | Helicase activity. |
| DEHA2E01232g | Superoxide metabolic process |
| DEHA2E03960g | Transient receptor potential (TRP) ion channel |
| DEHA2E05544g | Alkyl hydroperoxide reductase, thiol specific antioxidant and related enzymes |
| DEHA2E07546g | Flavin-containing monooxygenase |
| DEHA2E07590g | Voltage-dependent anion channel |
| DEHA2E07612g | C4-dicarboxylate transporter/malic acid transport protein |
| DEHA2E07634g | C4-dicarboxylate transporter/malic acid transport protein |
| DEHA2E11022g | Putative manganese resistance, Mg2+ transport protein MnR2p |
| DEHA2E15070g | The vacuolar basic amino acid (Arg, Lys, His) transporter, Vba3 |
| DEHA2F00550g | Ethionine resistance protein, ERC1 |
| DEHA2F01870g | The suppressor of sphingoid long chain base (LCB) sensitivity of an LCB-lyase mutation; putative flippase that exports LCBs across the cytoplasmic membrane, Rsb1p |
| DEHA2F03938g | Alcohol dehydrogenase, class V |
| DEHA2F11638g | Vacuolar H^+^-ATPase V0 sector, subunits c/c' |
| DEHA2F12650g | Permease of the major facilitator superfamily |
| DEHA2F14542g | Synaptic vesicle transporter SVOP and related transporters (major facilitator superfamily) |
| DEHA2F15642g | The multidrug efflux pump, Qdr3 (exports polyamines, quinidine, barban, cisplatin and bleomycin) |
| DEHA2F16434g | RNA-binding protein (contains RRM and Pumilio-like repeats) |
| DEHA2G02420g | Thioredoxin binding protein TBP-2/VDUP1 |
| DEHA2G05808g | Alcohol dehydrogenase, class III |
| DEHA2G09372g | Aldehyde dehydrogenase |
| DEHA2G13244g | Putative Zn^2+^ transporter MSC2 (cation diffusion facilitator superfamily) |
| DEHA2G14916g | Multidrug resistance protein, Cdr1 (Candida drug resistance 1) confers resistance to cycloheximide, xenobiotics and antifungal agents such as azoles and terbinafine. |
| DEHA2B06974g | Multicopper oxidases |
| DEHA2C01562g | Aldehyde dehydrogenase |
| DEHA2E05940g | Acetylcholinesterase/Butyrylcholinesterase/Carboxylesterase family |
| DEHA2G09372g | Oxidoreductase activity. Aldehyde dehydrogenase family |
| **ORFs involved in glutathione metabolism** | |
| DEHA2A02706g | Oxoprolinase |
| DEHA2C10274g | Glucose-6-phosphate 1-dehydrogenase |
| DEHA2D06160g | 6-phosphogluconate dehydrogenase |
| DEHA2D09130g | Gamma-glutamyltransferase |
| DEHA2D16280g | Glutathione S-transferase |
| DEHA2D16302g | Glutathione S-transferase |
| DEHA2E10692g | NADP-dependent isocitrate dehydrogenase |
| DEHA2G04312g | Hydantoinase/Oxoprolinase |
| DEHA2G19844g | Metalloexopeptidases |
| **ORFs with oxidoreductase activity, which acts on the CH-OH group** | |
| DEHA2A06116g | 6-phosphogluconate dehydrogenase, NAD-binding |
| DEHA2B03058g | NAD-dependent malate dehydrogenase |
| DEHA2B07678g | Ketopantoate reductase |
| DEHA2C08602g | Flavonol reductase/cinnamoyl-CoA reductase |
| DEHA2C10758g | Isocitrate dehydrogenase, gamma subunit |
| DEHA2C11550g | C-3 sterol dehydrogenase/3-beta-hydroxysteroid dehydrogenase and related dehydrogenases |
| DEHA2C16852g | Isocitrate dehydrogenase, alpha subunit |
| DEHA2D06160g | 6-phosphogluconate dehydrogenase |
| DEHA2E10692g | NADP-dependent isocitrate dehydrogenase |
| DEHA2F09020g | NAD-dependent malate dehydrogenase |
| DEHA2F09570g | Acetohydroxy acid isomeroreductase |
| DEHA2F10274g | Nuclear receptor coregulator SMRT/SMRTER, contains Myb-like domains |
| DEHA2G05786g | Isocitrate dehydrogenase, alpha subunit |
| DEHA2G19360g | Glyoxylate/hydroxypyruvate reductase (D-isomer-specific 2-hydroxy acid dehydrogenase superfamily) |
| **Methane metabolism** | |
| DEHA2A03542g | Dihydroxyacetone kinase/glycerone kinase |
| DEHA2D02112g | Enolase |
| DEHA2D10186g | Pyrophosphate-dependent phosphofructo-1-kinase |
| DEHA2E01694g | Alanine-glyoxylate aminotransferase AGT1 |
| DEHA2E04708g | Esterase D |
| DEHA2E05676g | Acyl-CoA synthetase |
| DEHA2E06600g | Glycine/serine hydroxymethyltransferase |
| DEHA2G05808g | Alcohol dehydrogenase, class III |
| DEHA2G14058g | Enolase |
| DEHA2G19360g | Glyoxylate/hydroxypyruvate reductase (D-isomer-specific 2-hydroxy acid dehydrogenase superfamily) |
| DEHA2G20768g | Pyrophosphate-dependent phosphofructo-1-kinase |
| **Propionate metabolism** | |
| DEHA2A06314g | Flavonol reductase/cinnamoyl-CoA reductase |
| DEHA2B06424g | Predicted acyl-CoA dehydrogenase |
| DEHA2C16324g | 4-aminobutyrate aminotransferase |
| DEHA2E05676g | Acyl-CoA synthetase |
| DEHA2F09306g | 4-aminobutyrate aminotransferase |
| DEHA2G12342g | Enoyl-CoA hydratase |
| **Butanoate metabolism** | |
| DEHA2C16324g | 4-aminobutyrate aminotransferase |
| DEHA2F09306g | 4-aminobutyrate aminotransferase |
| DEHA2F10450g | Glutamate decarboxylase/sphingosine phosphate lyase |
| DEHA2F16896g | Sorbitol dehydrogenase |
| DEHA2G02134g | Acetolactate synthase, large subunit, biosynthetic |
| **Catabolism of C-2 compounds and organic acids** | |
| DEHA2C00880g | Alcohol dehydrogenase, class III |
| DEHA2D10120g | Thiamine pyrophosphate-requiring enzyme |
| DEHA2D12936g | Isocitrate lyase |
| DEHA2D16984g | Thiamine pyrophosphate-requiring enzyme |
| DEHA2E05676g | Acyl-CoA synthetase |
| DEHA2F00792g | Aldehyde dehydrogenase |
| DEHA2F03938g | Alcohol dehydrogenase, class V |
| DEHA2G18348g | Thiamine pyrophosphate-requiring enzyme |
| DEHA2G23650g | C1-tetrahydrofolate synthase |
| **Glyoxylate and dicarboxylate metabolism** | |
| DEHA2B03058g | NAD-dependent malate dehydrogenase |
| DEHA2C01584g | Isocitrate lyase |
| DEHA2D12936g | Isocitrate lyase and phosphorylmutase |
| DEHA2E01694g | Alanine-glyoxylate aminotransferase AGT1 |
| DEHA2E06600g | Glycine/serine hydroxymethyltransferase |
| DEHA2E13530g | Malate synthase |
| DEHA2F09020g | NAD-dependent malate dehydrogenase |
| DEHA2F20394g | Citrate synthase |
| DEHA2G06952g | Predicted kinase |
| DEHA2G19360g | Glyoxylate/hydroxypyruvate reductase (D-isomer-specific 2-hydroxy acid dehydrogenase superfamily) |
