## Supplementary figures and images for "Transcriptomic profiling of *Debaryomyces hansenii* reveals detoxification and stress responses to benzo(a)pyrene exposure"

### Supplementary_Figure_1.pptx

#### Slide 1
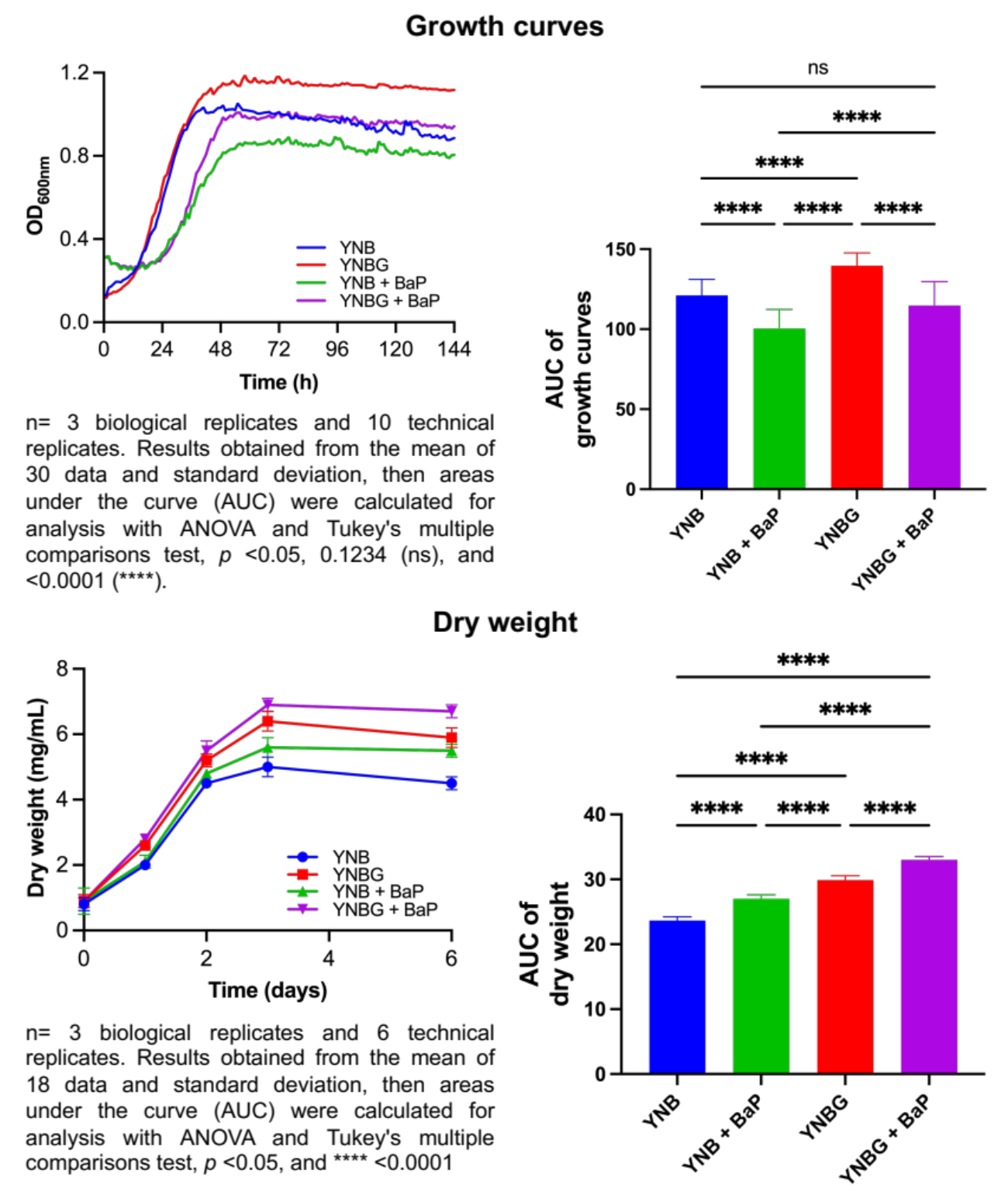

#### Slide 2
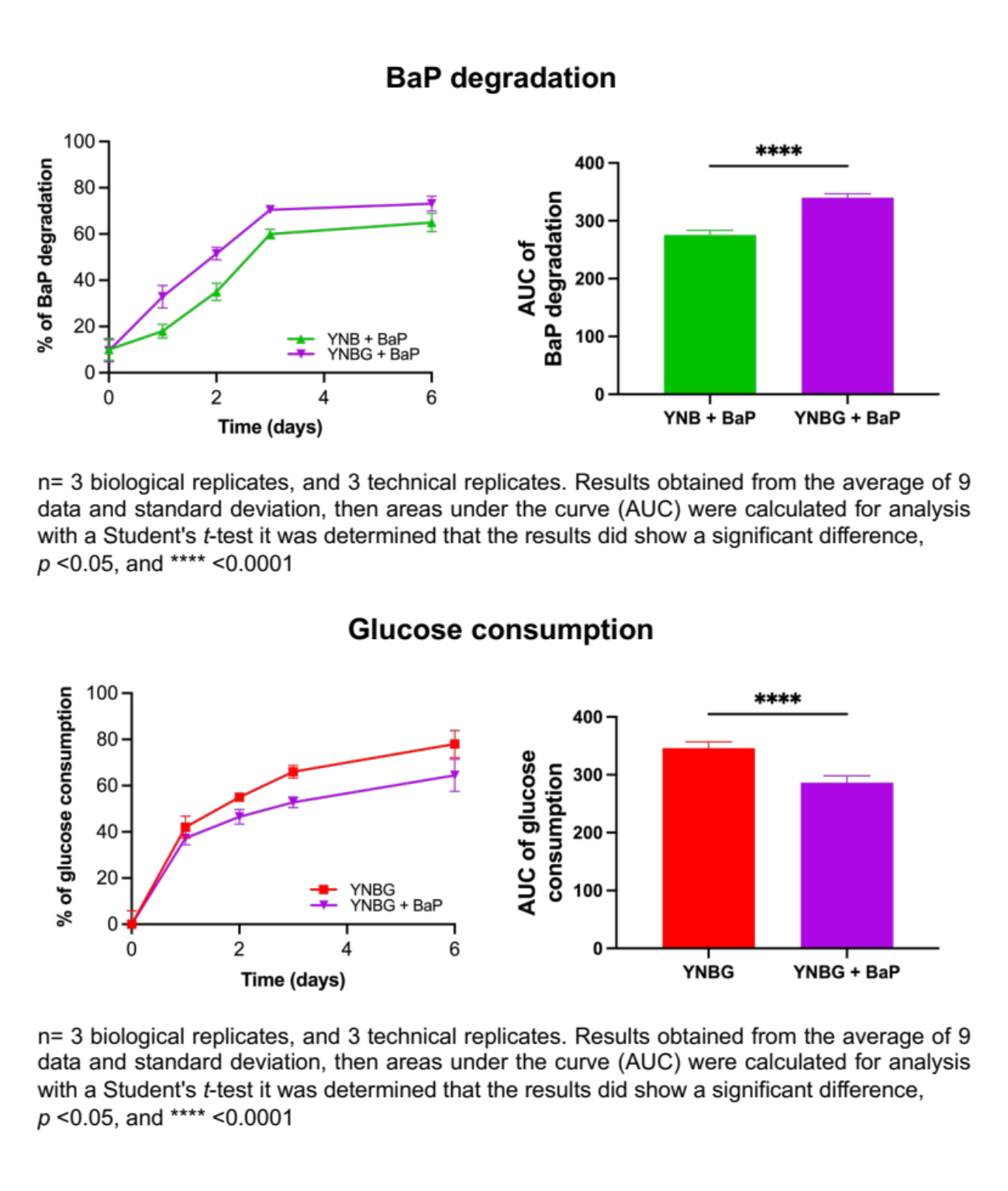

### Supplementary_Figure_2.pptx

#### Slide 1
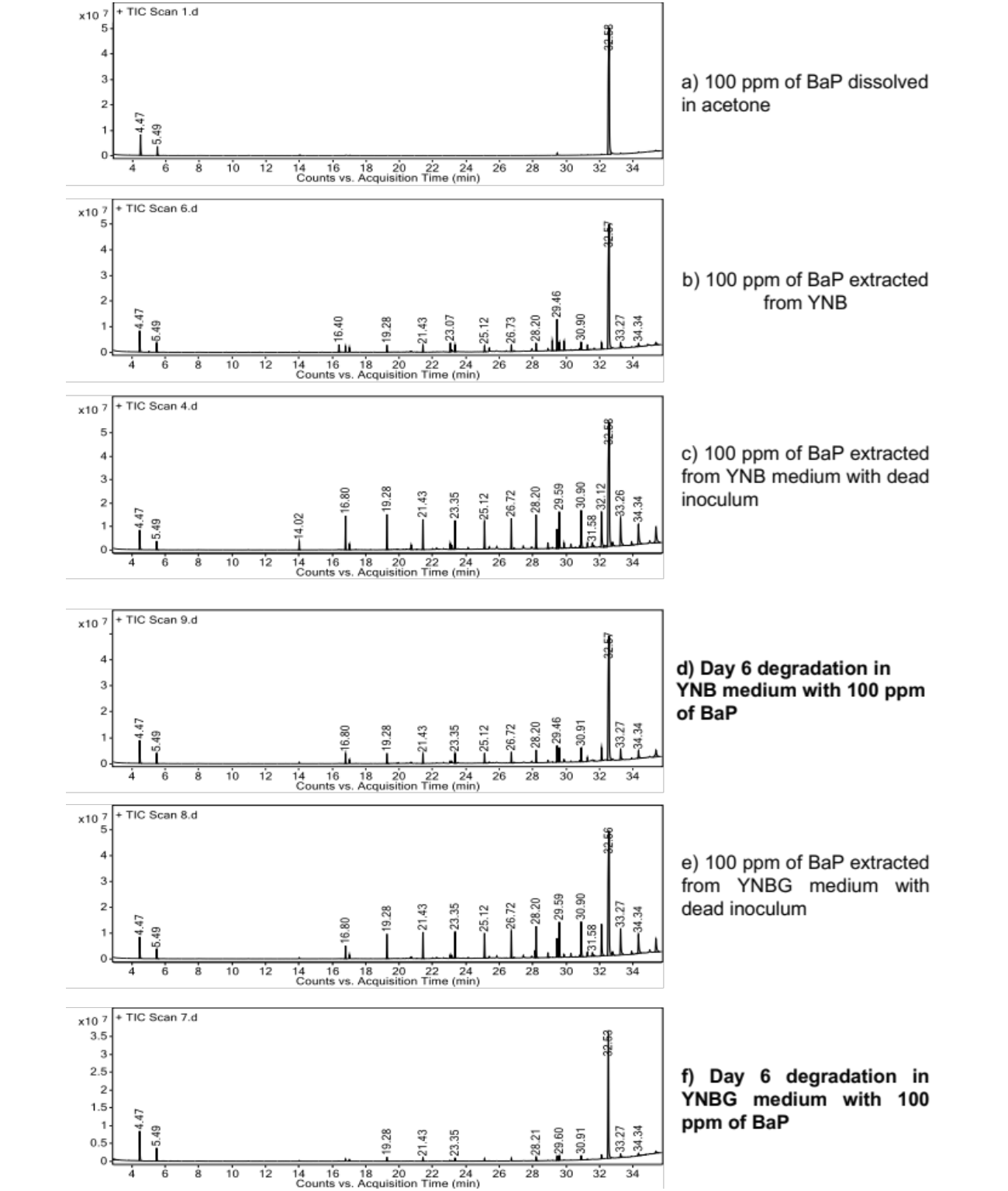

### Supplementary_Figure_3.pptx

#### Slide 1
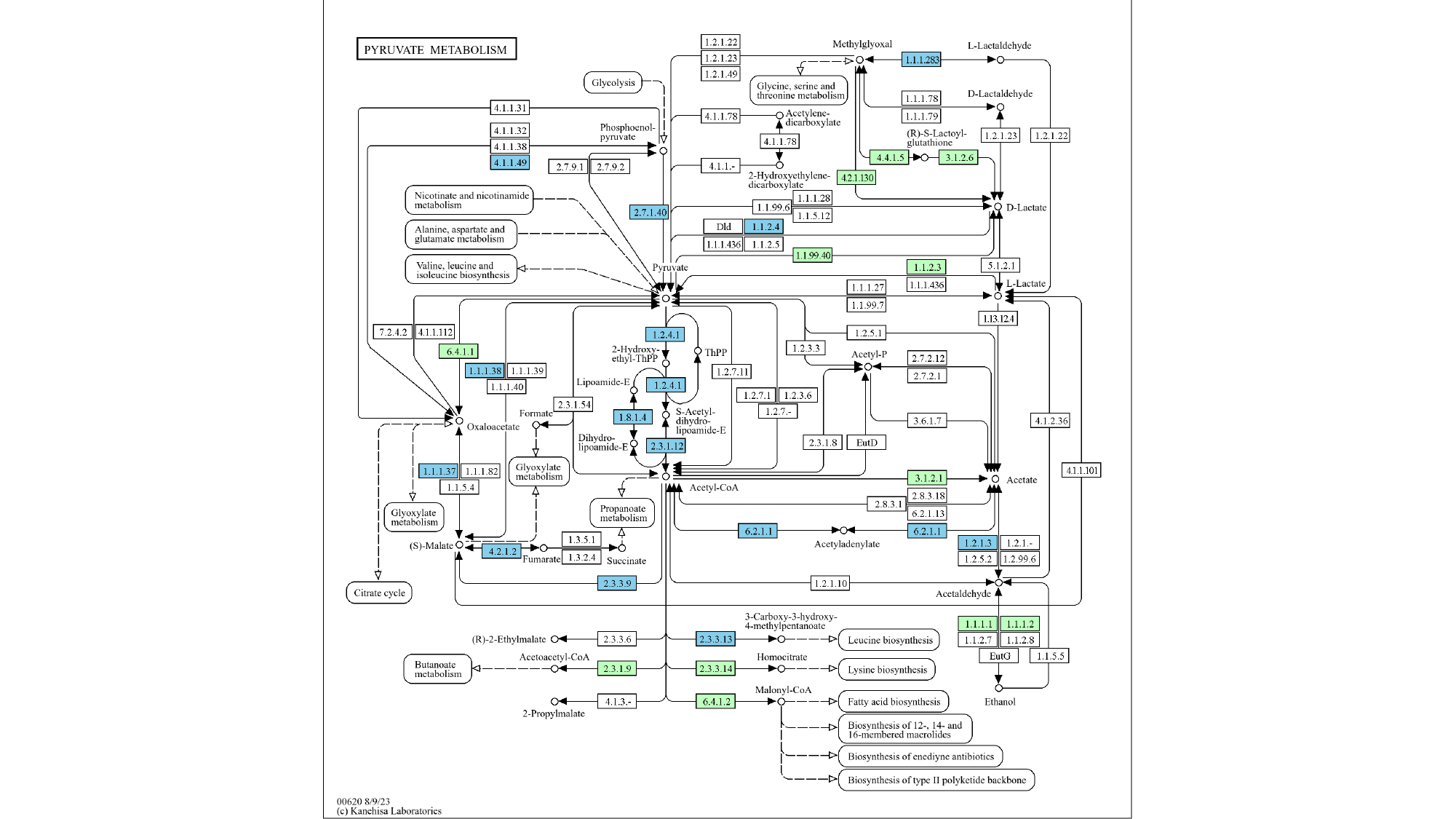

#### Slide 2
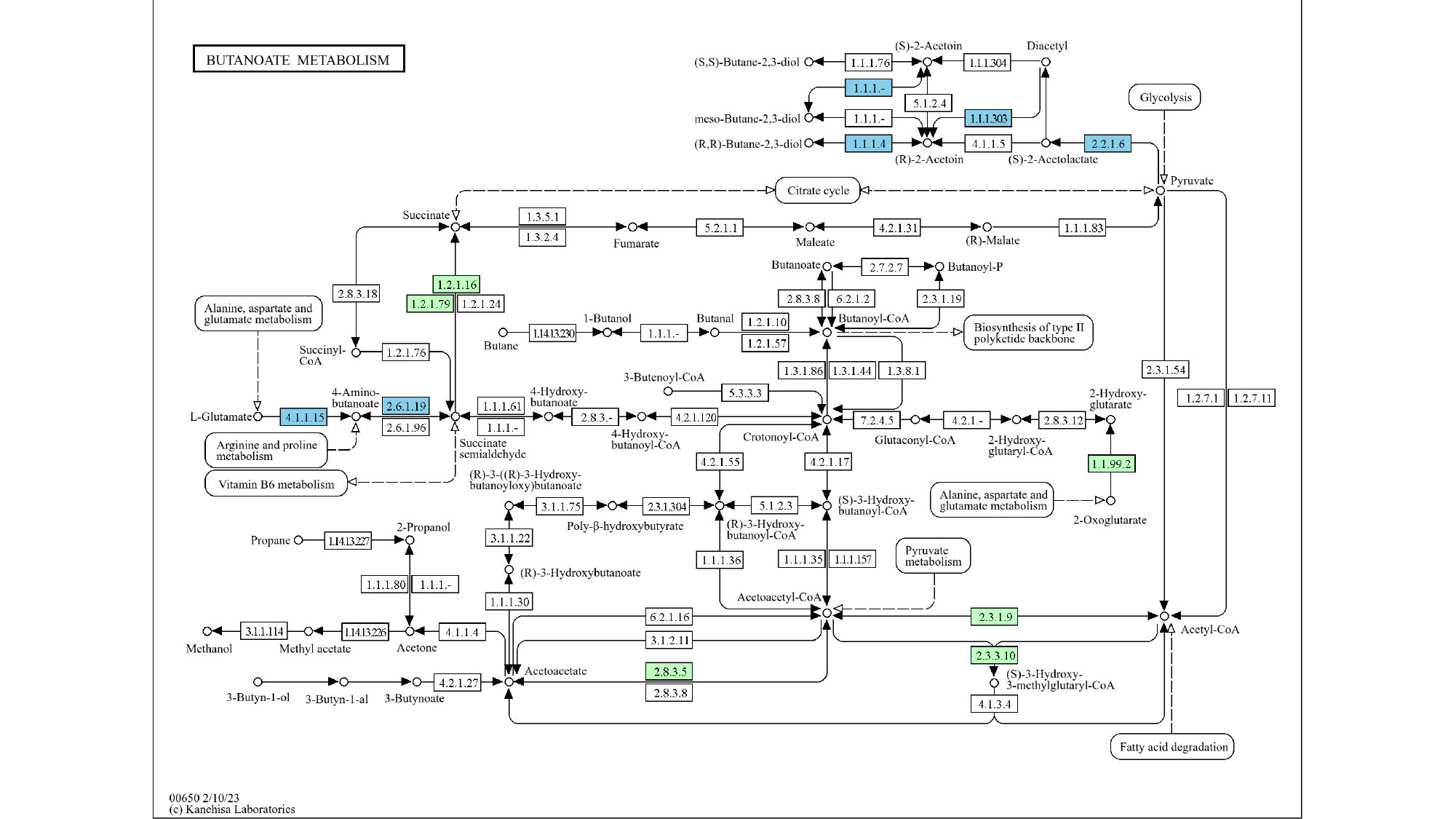

#### Slide 3
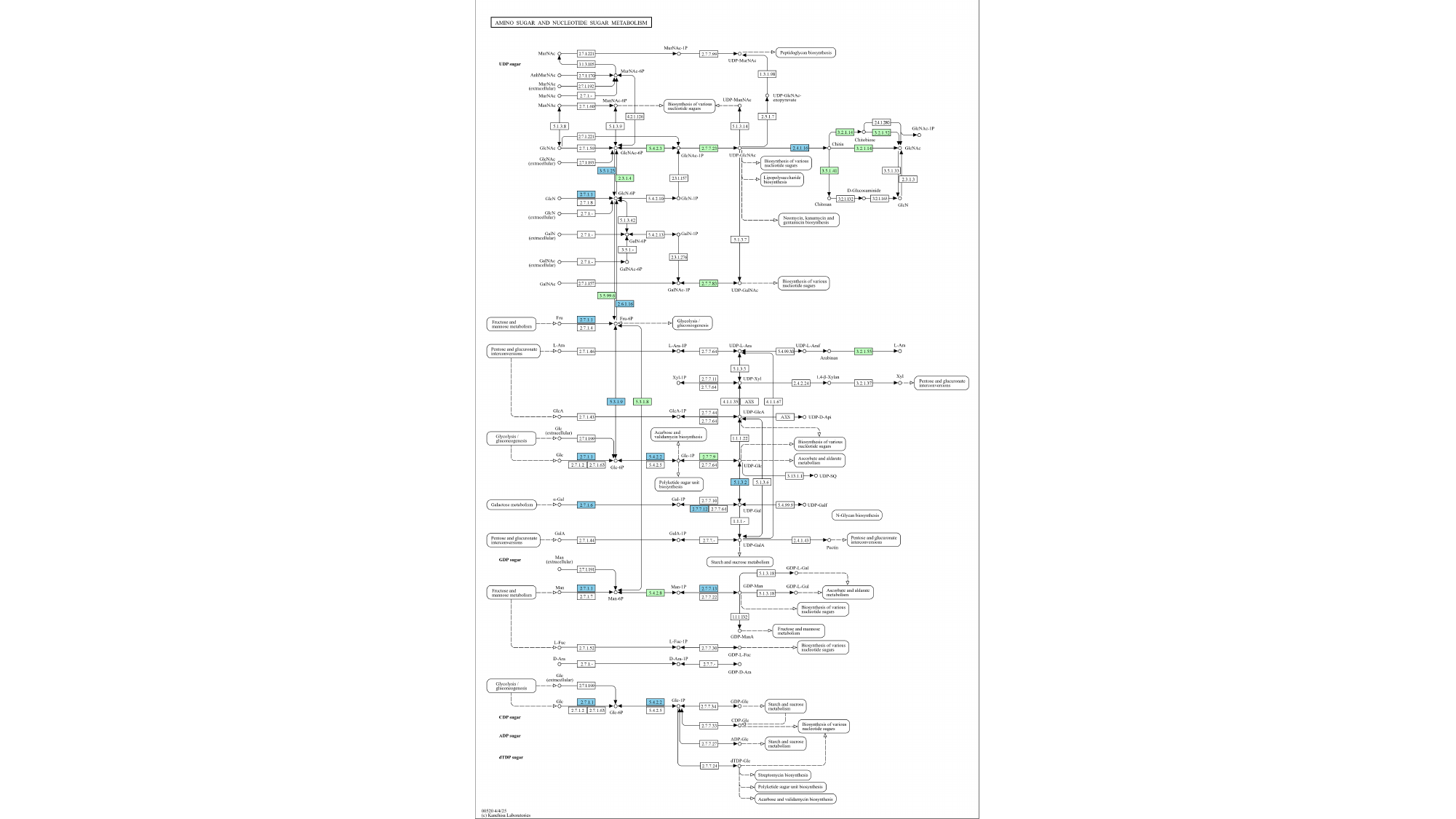

#### Slide 4
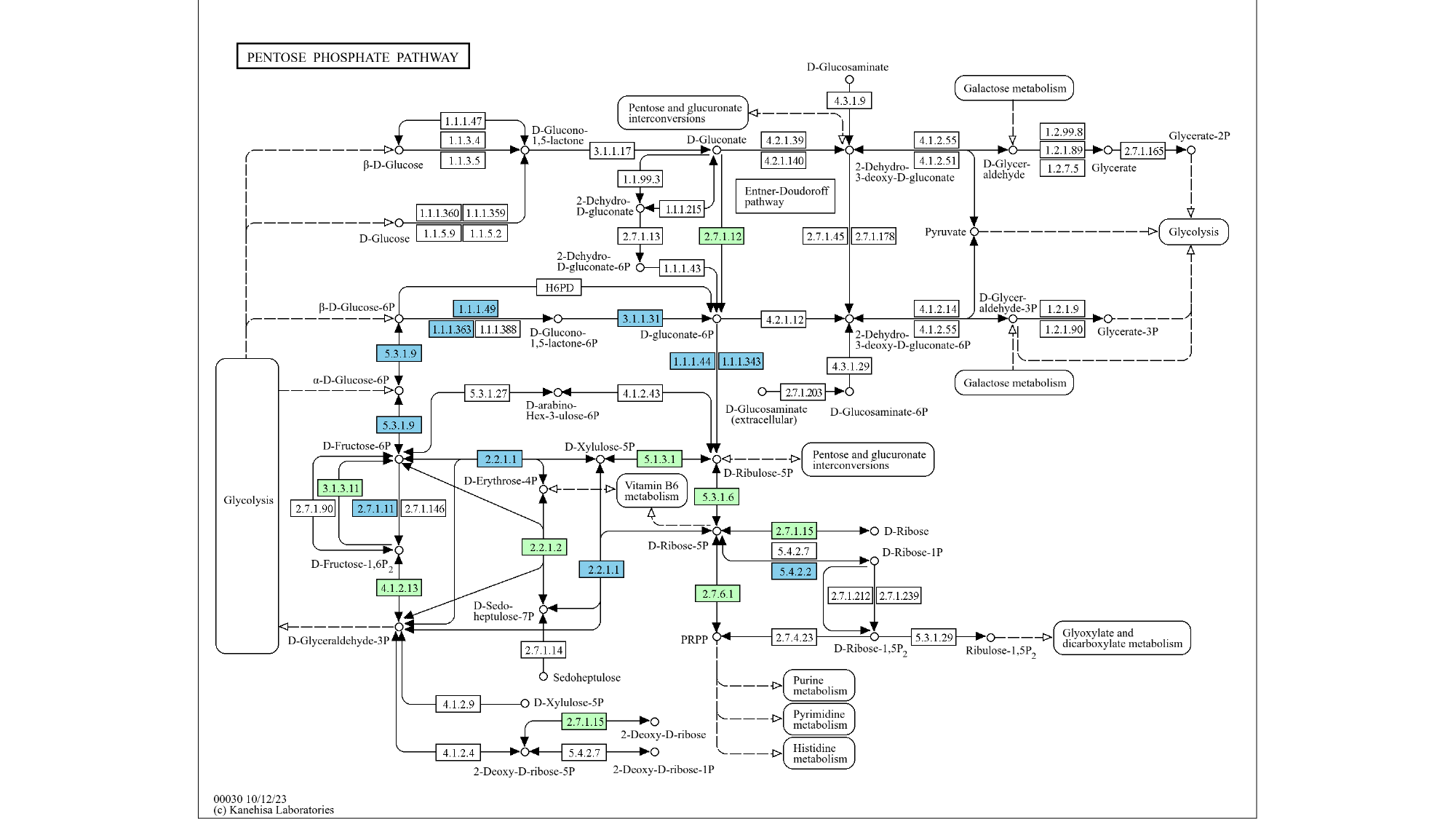

#### Slide 5
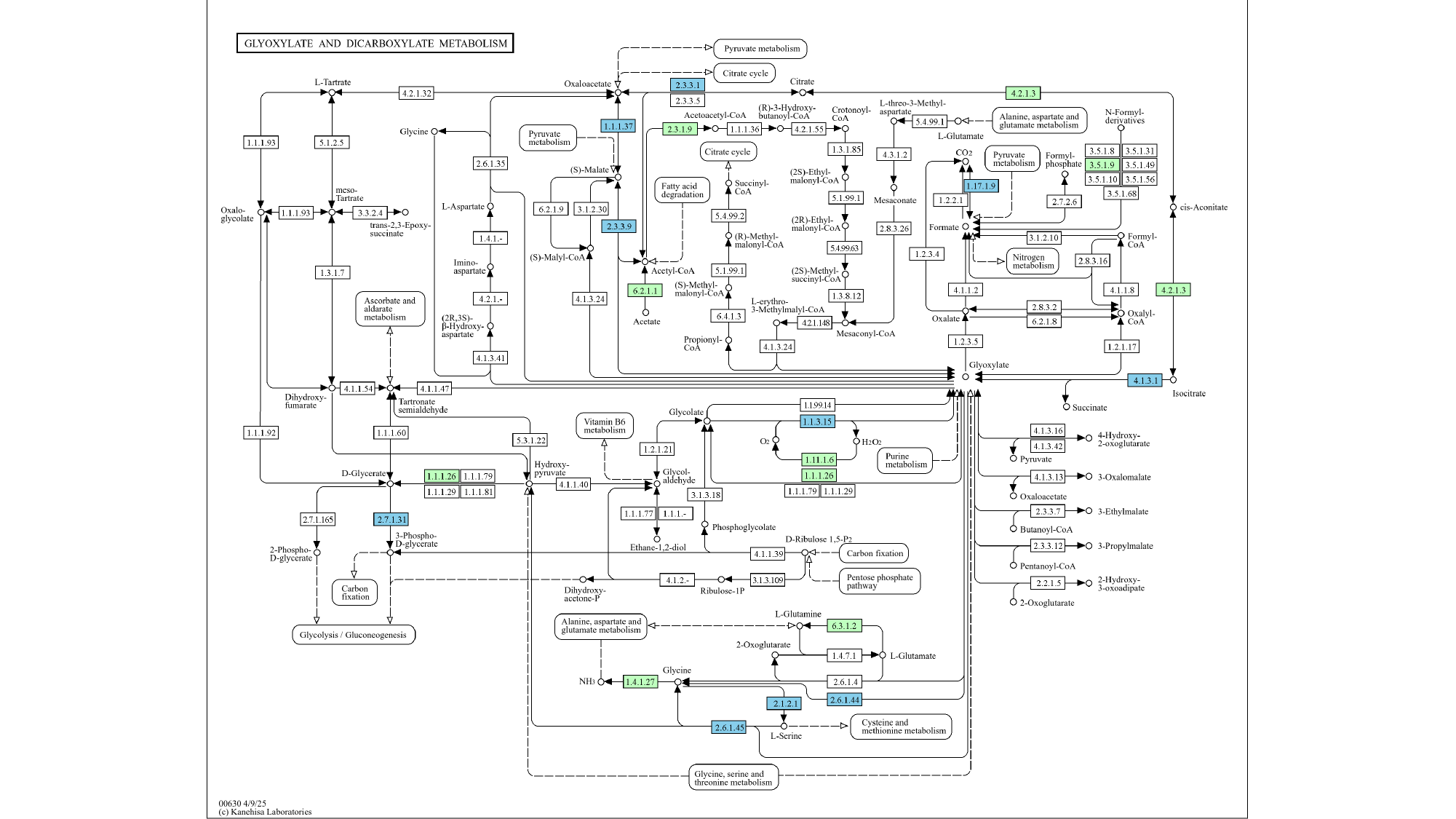

#### Slide 6
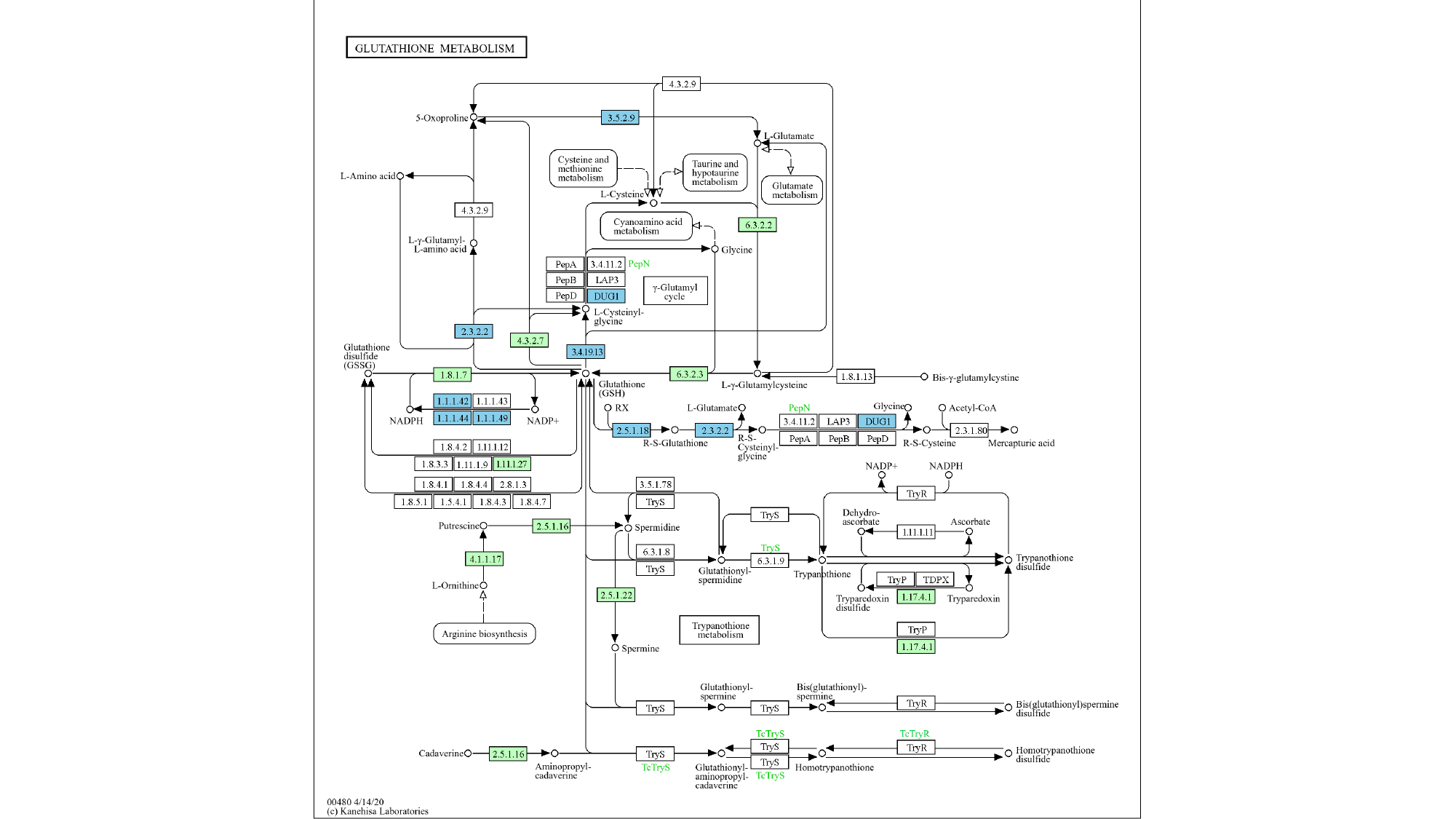

#### Slide 7
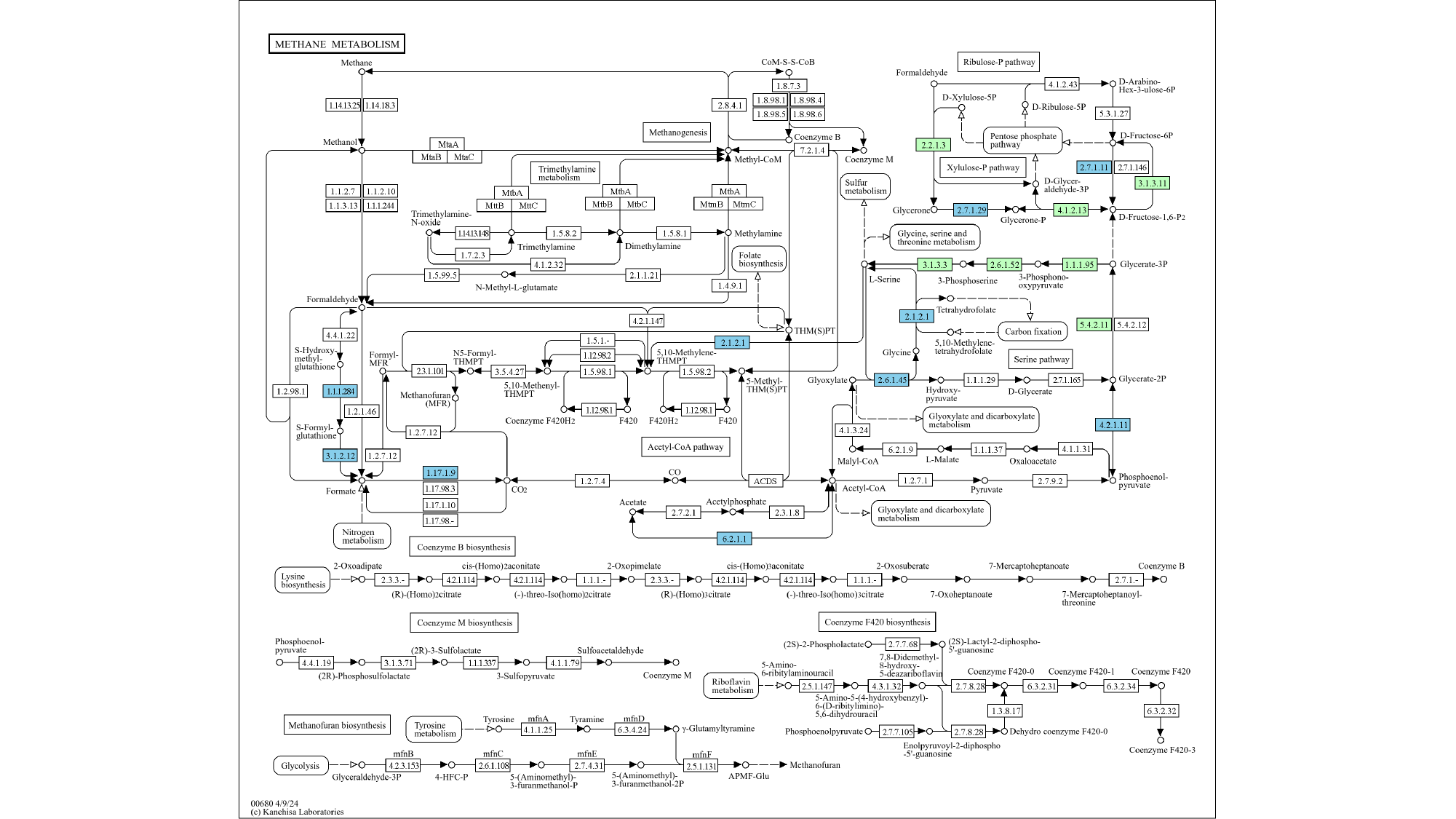

#### Slide 8
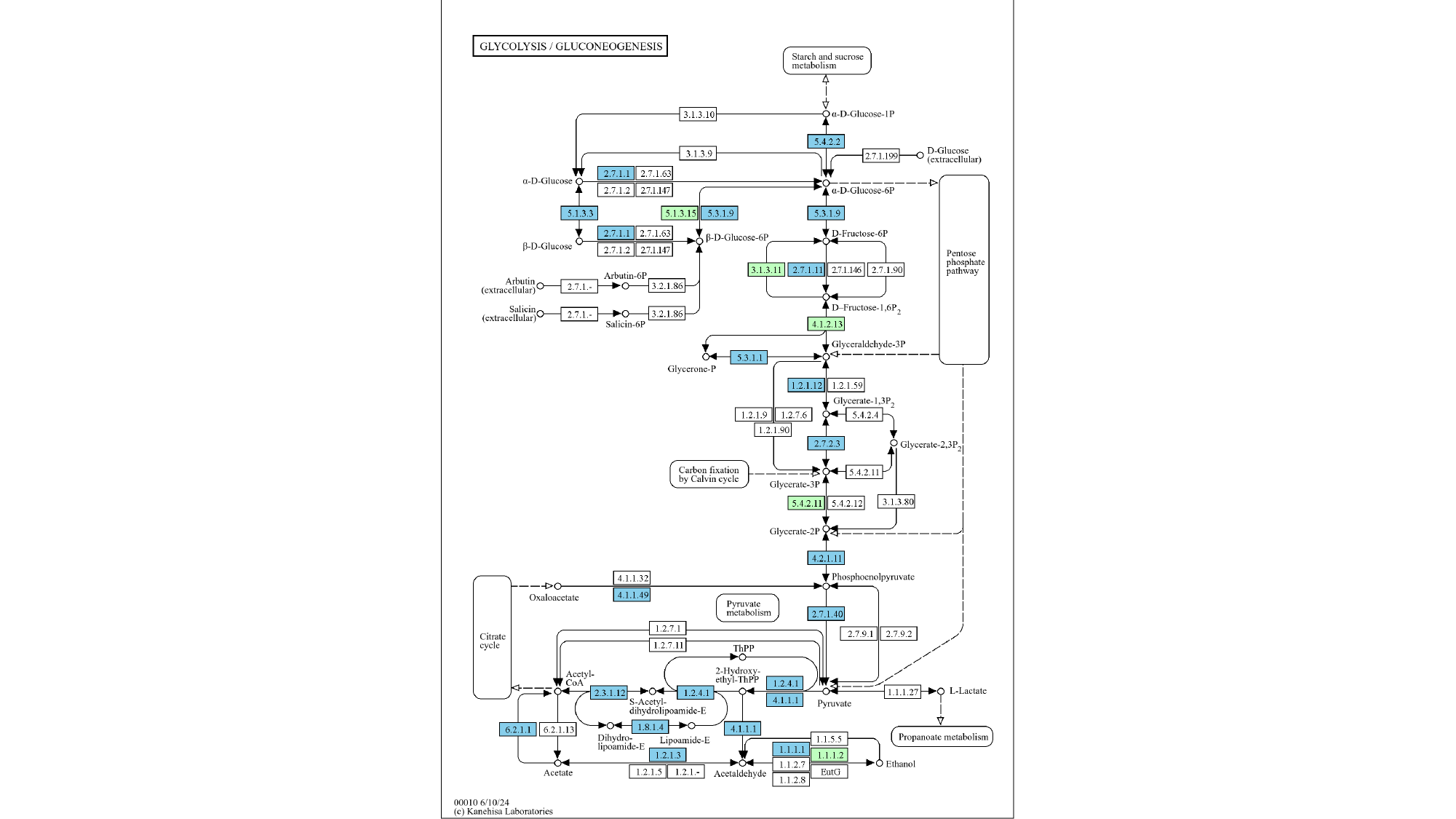

#### Slide 9
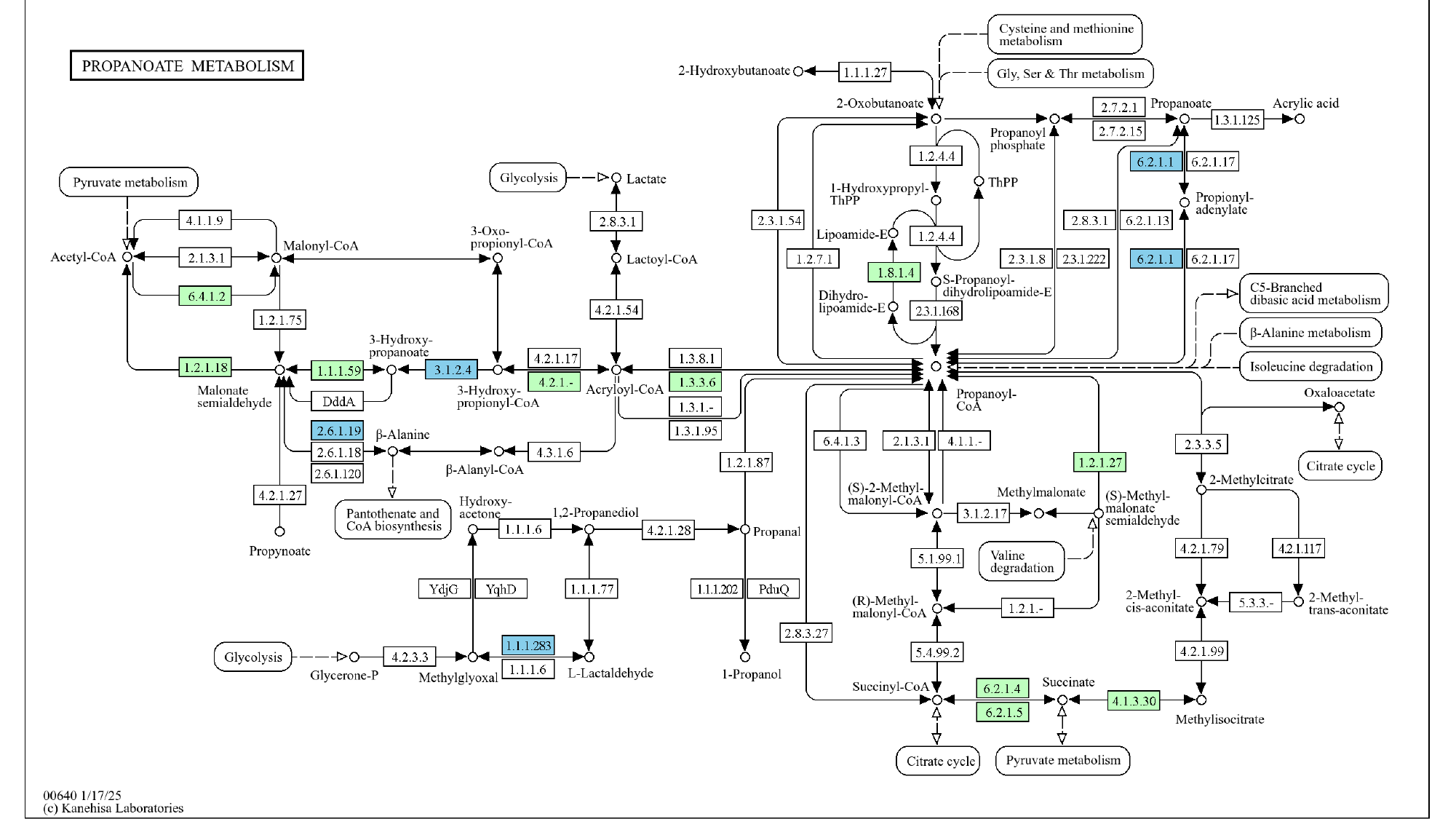

#### Slide 10
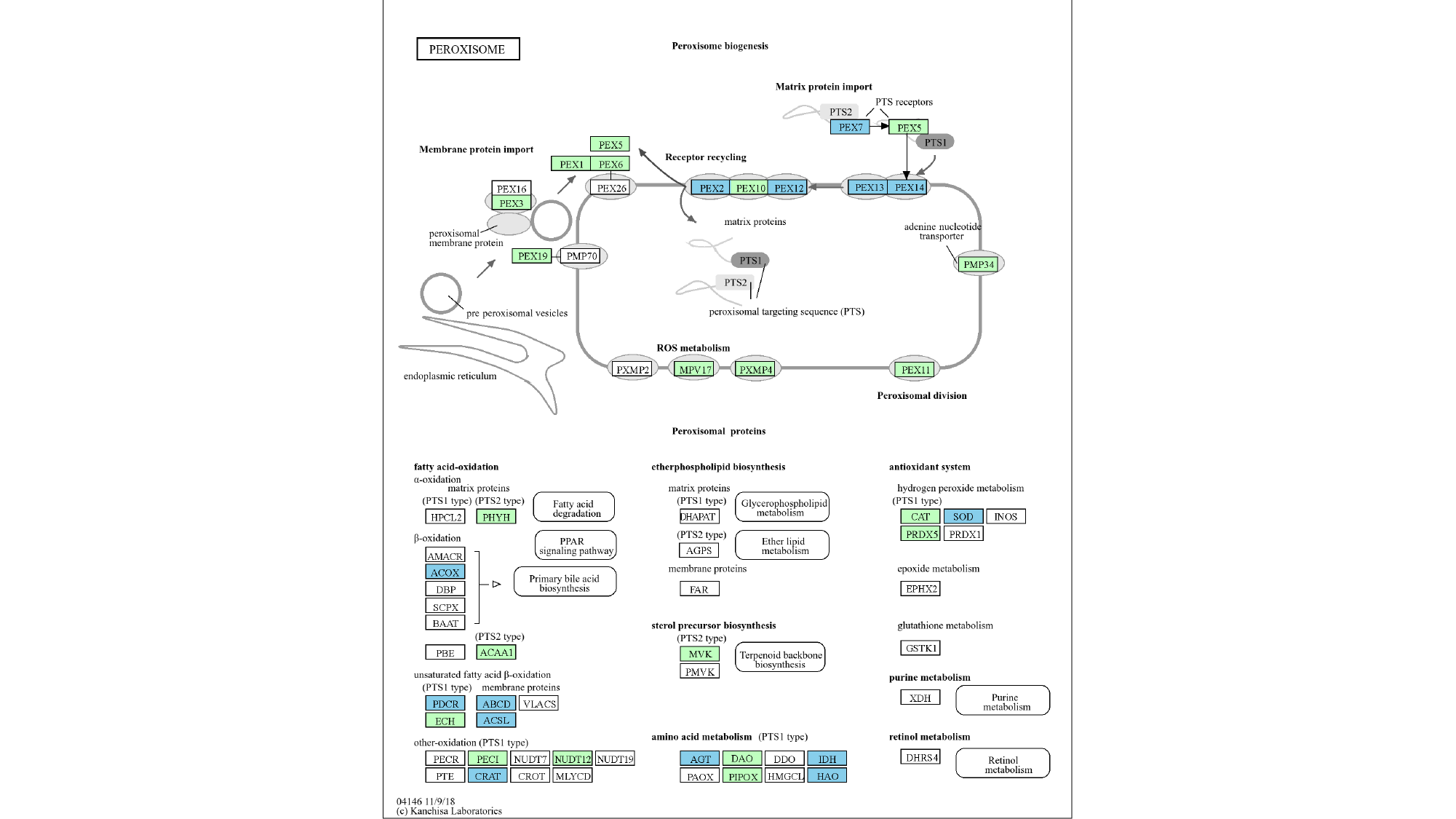

#### Slide 11
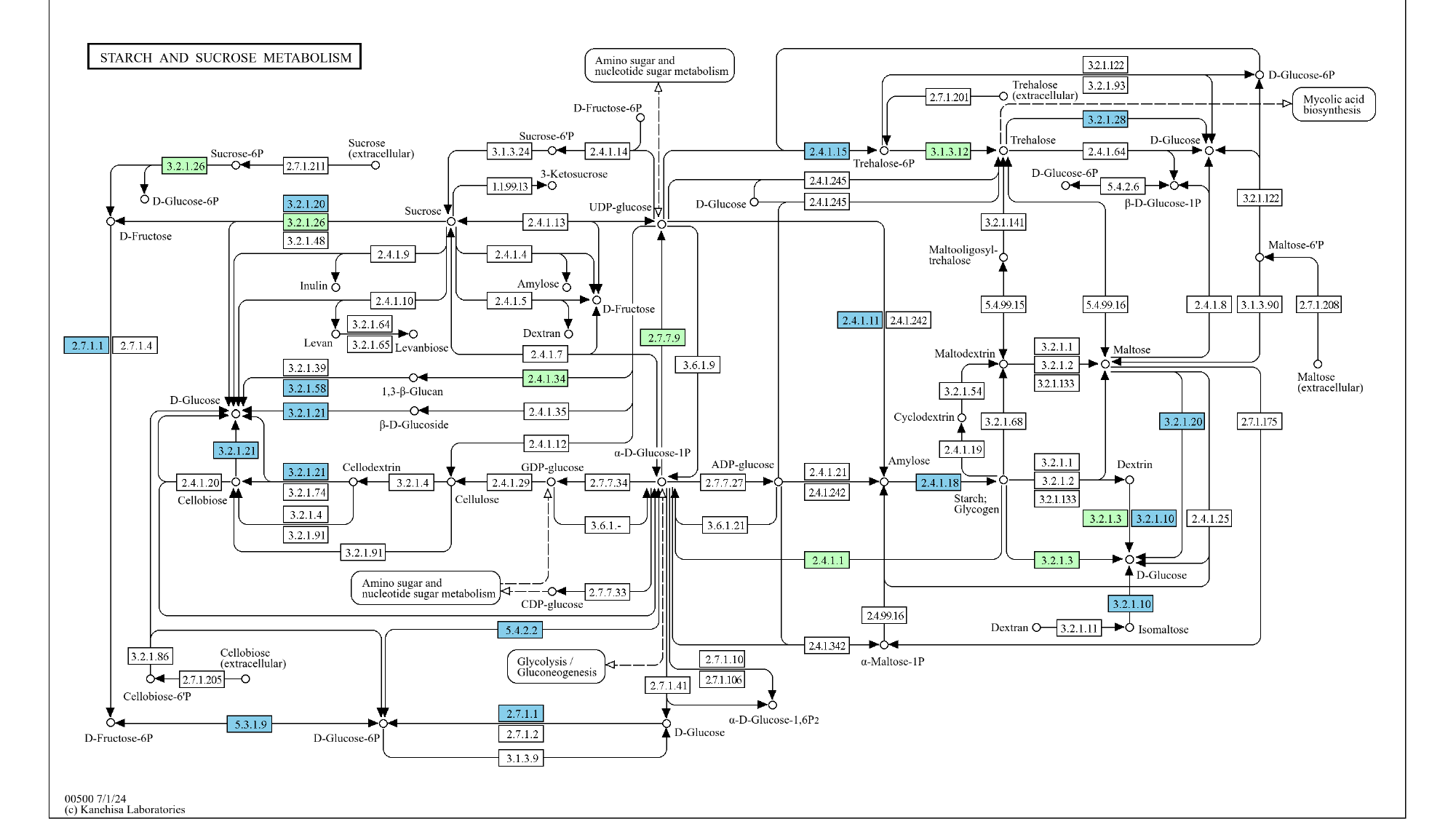

#### Slide 12
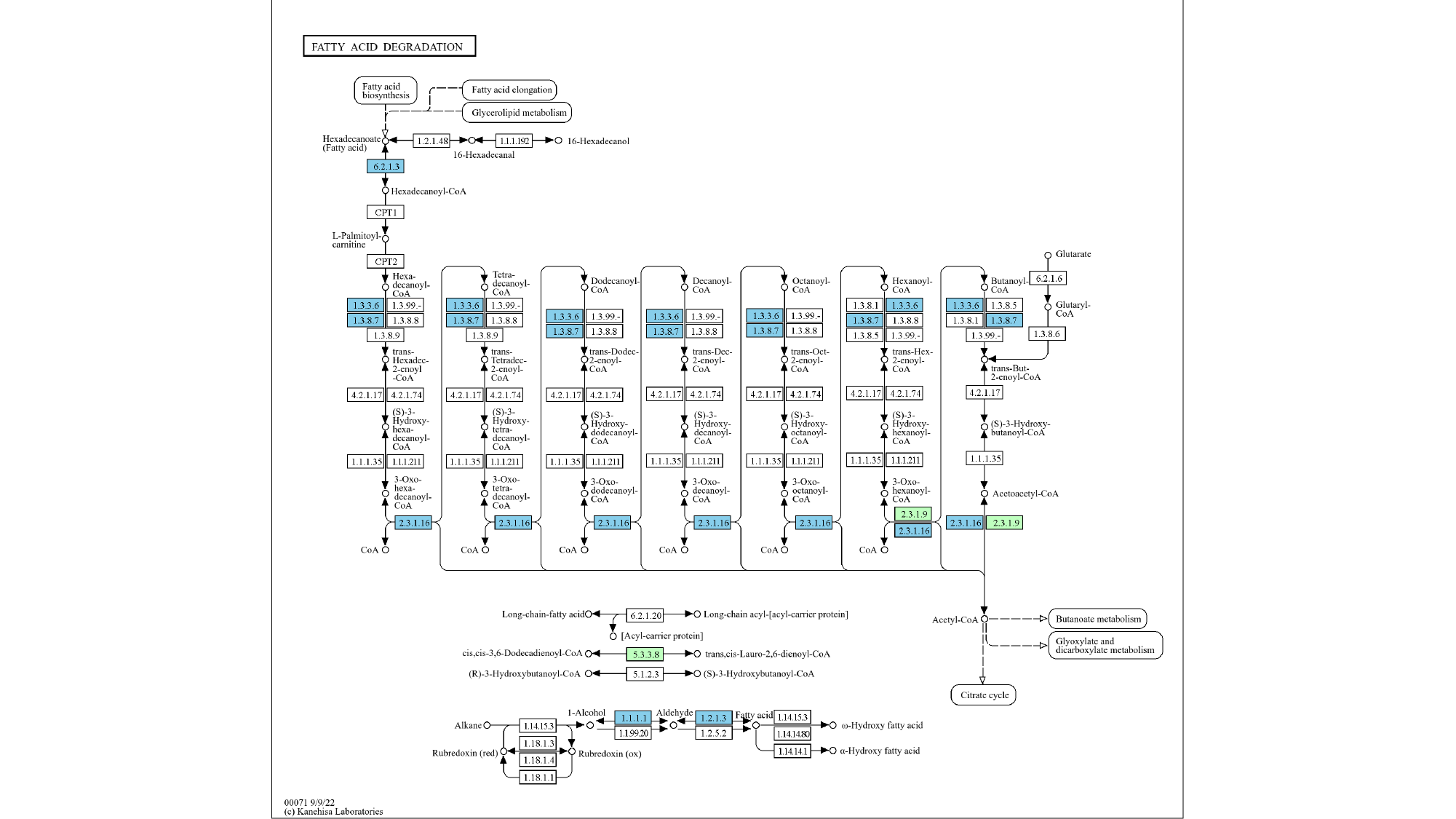

#### Slide 13
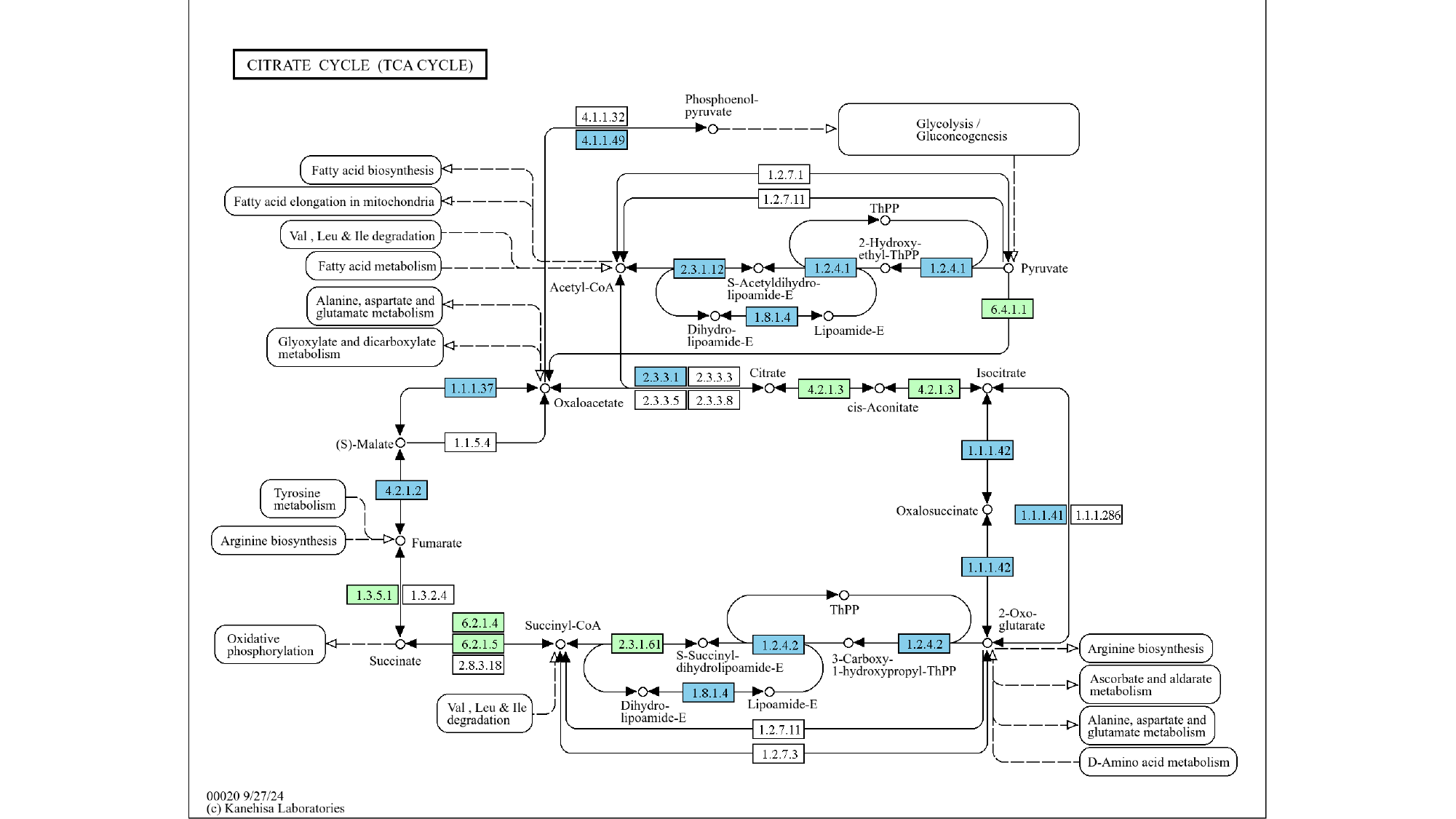

### Supplementary_Figure_4.pptx

#### Slide 1
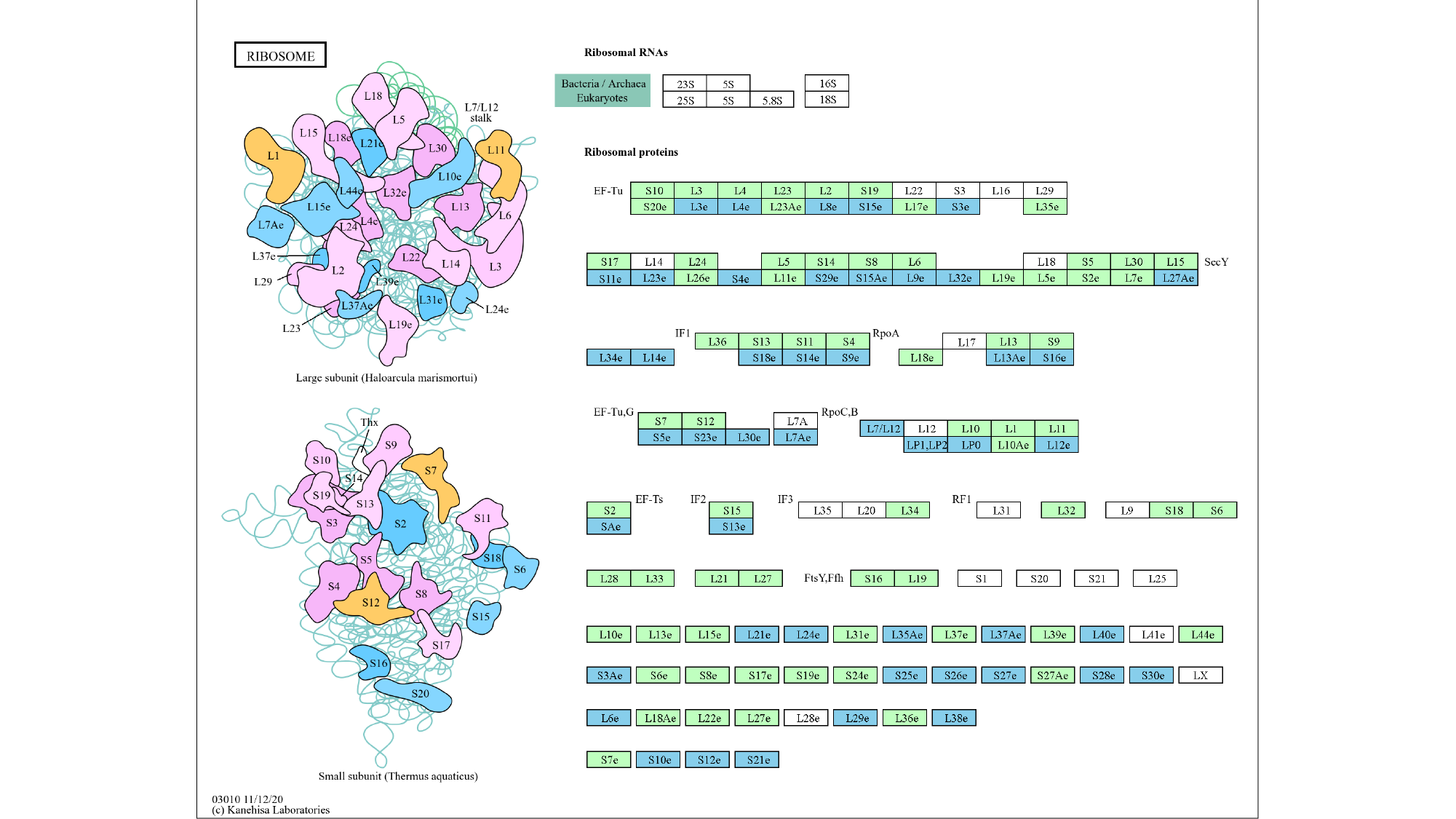

#### Slide 2
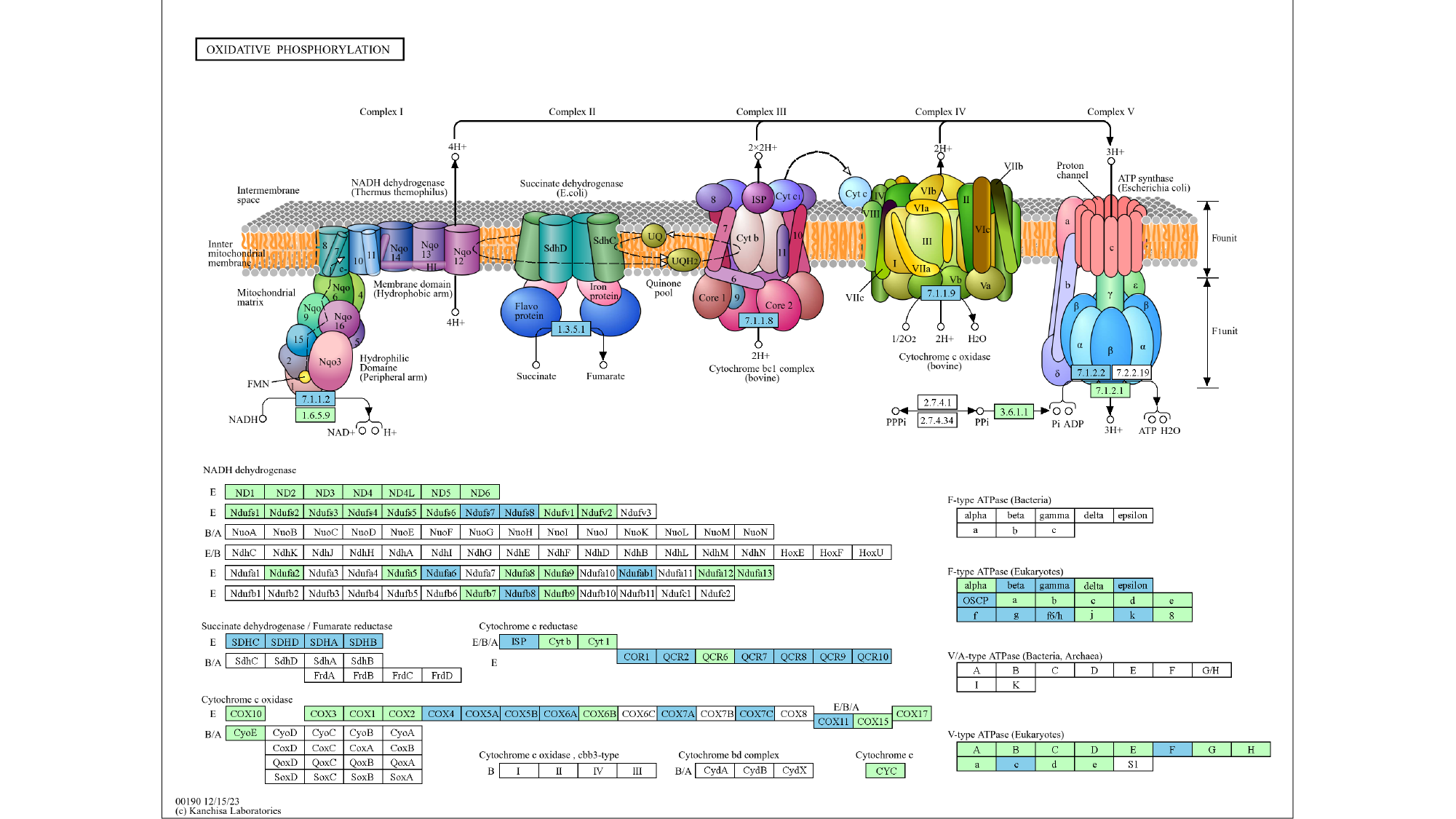

#### Slide 3
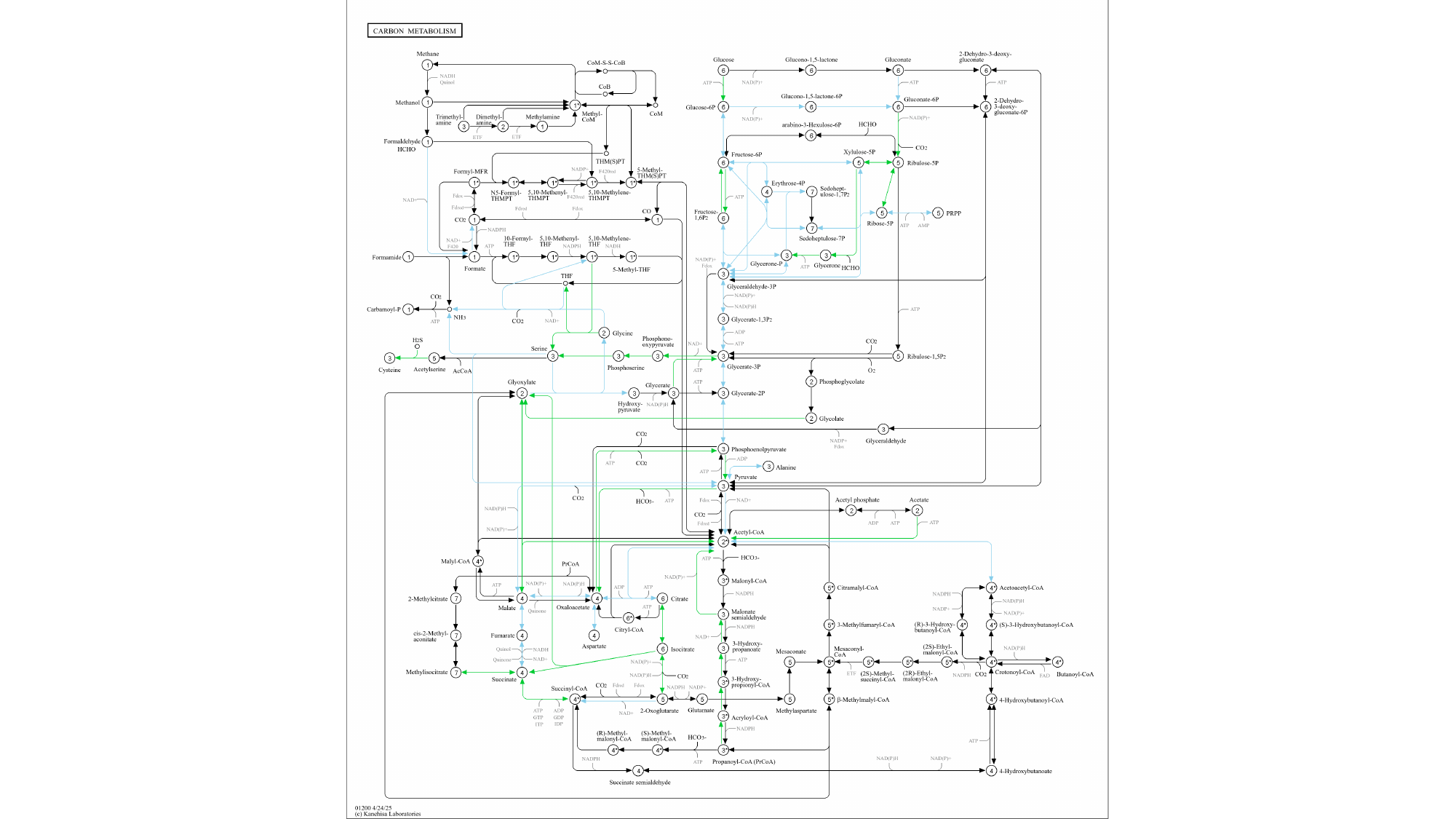

#### Slide 4
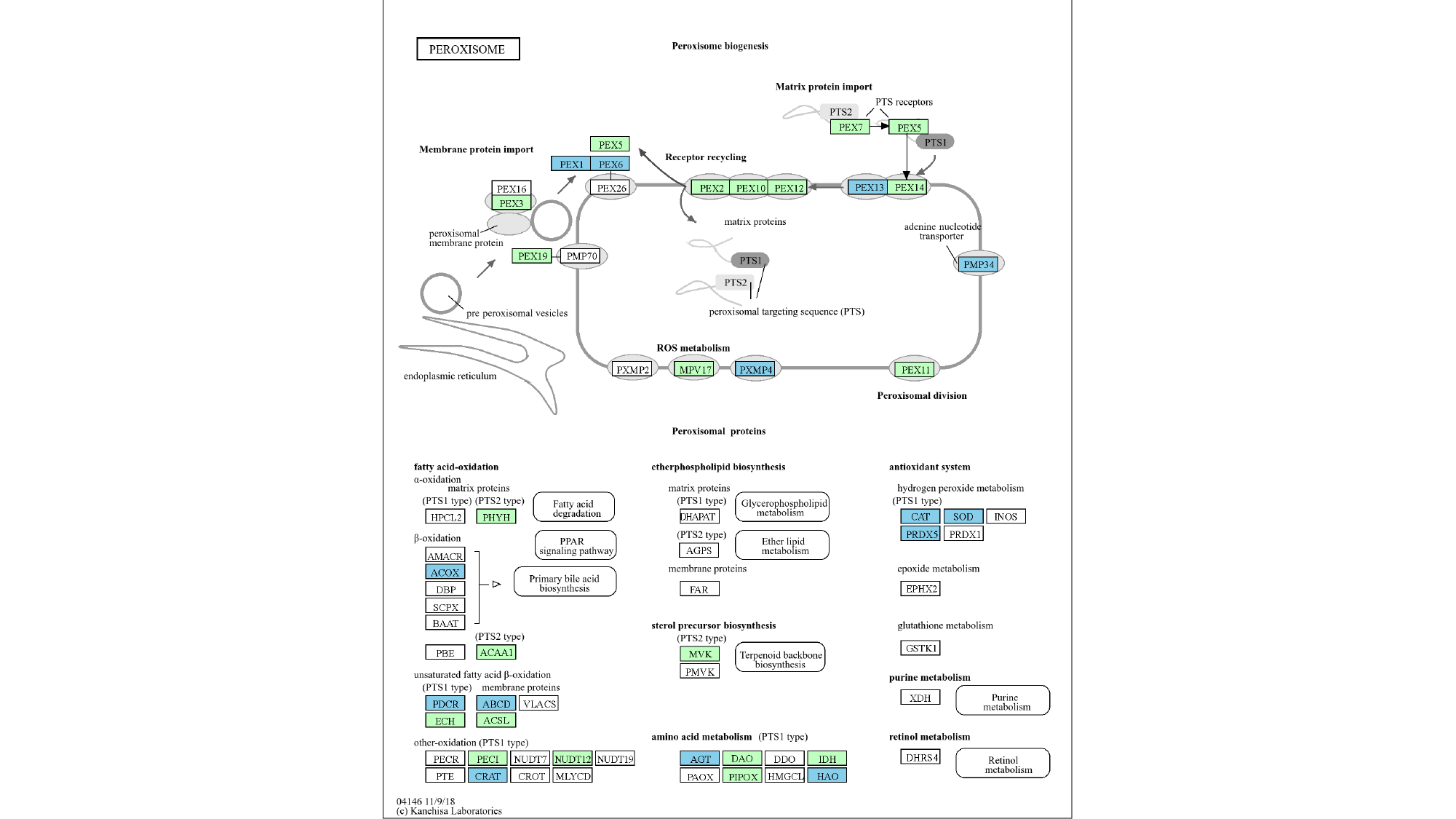

#### Slide 5
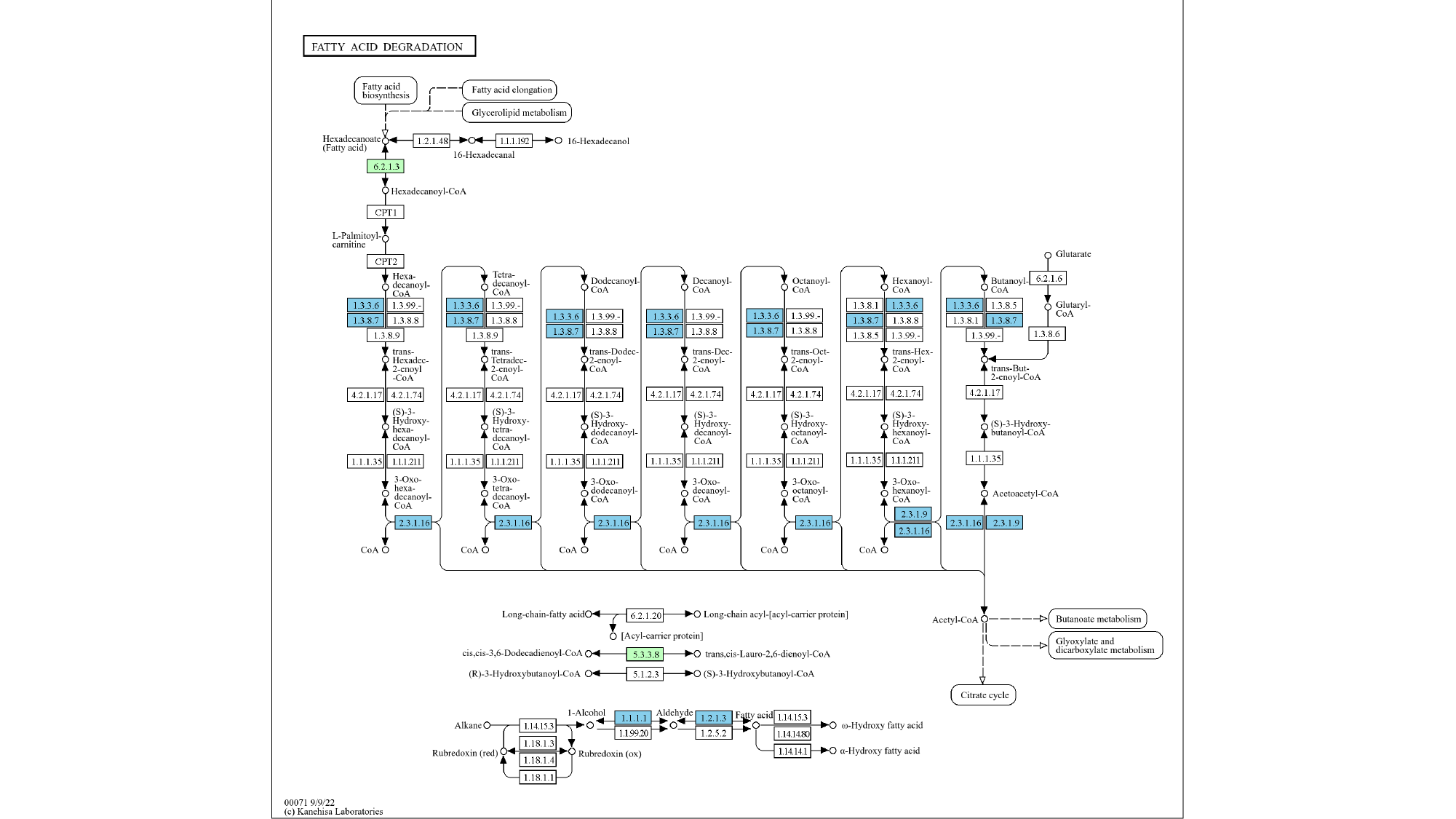

#### Slide 6
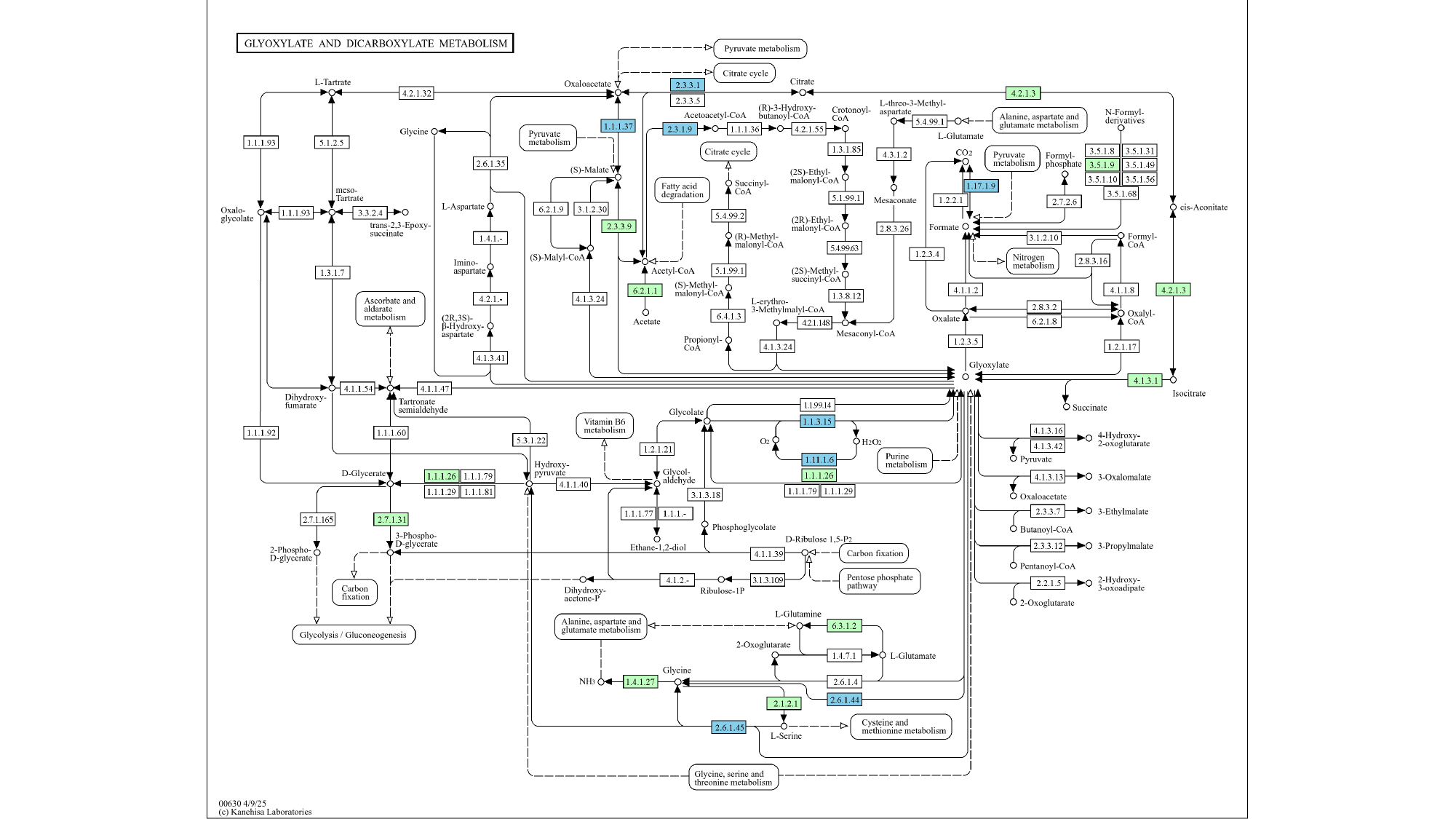

#### Slide 7
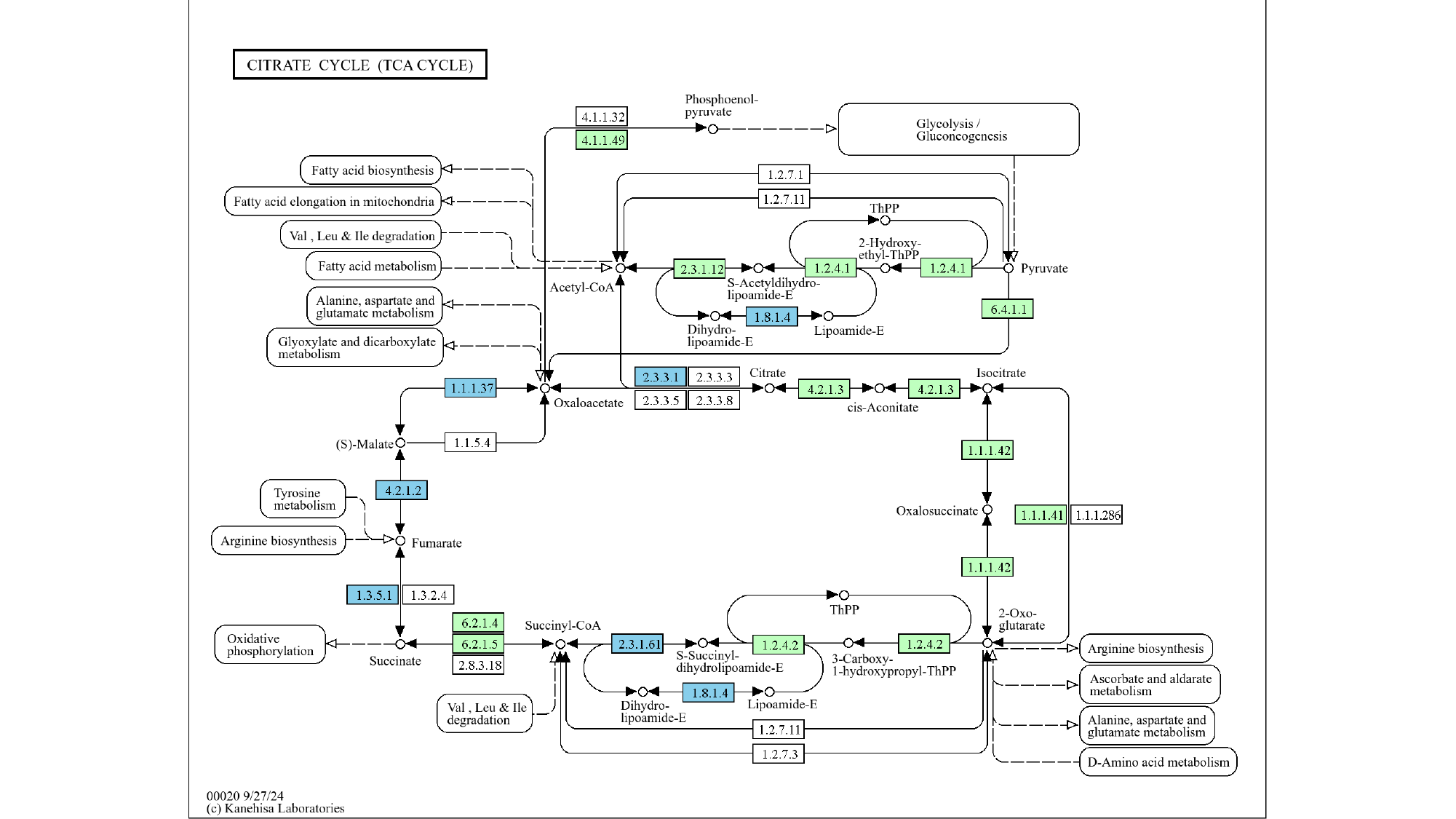

#### Slide 8
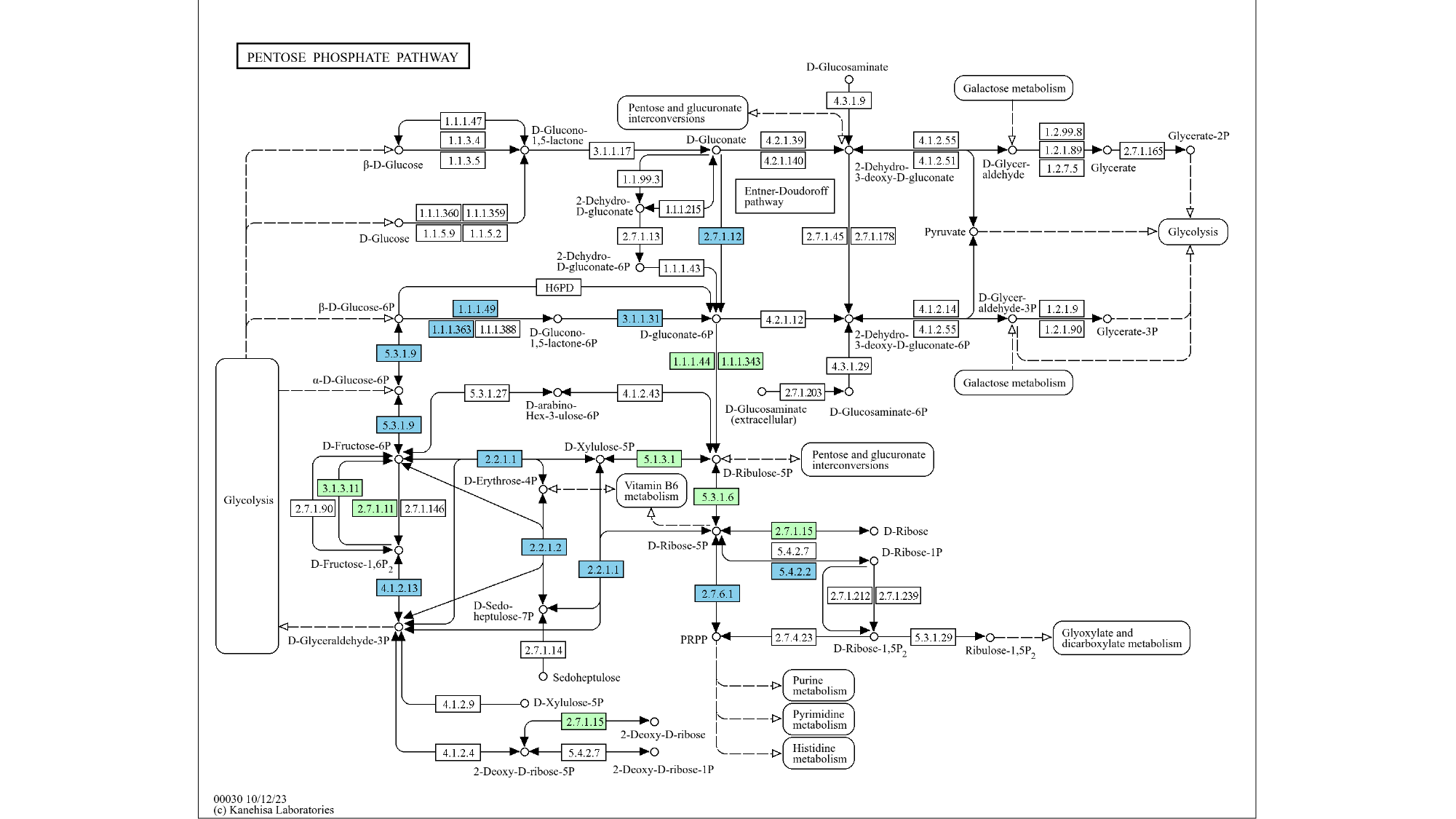
